# Large Mammals Have More Powerful Antibacterial Defenses Than Expected from Their Metabolic Rates

**DOI:** 10.1101/2020.09.04.242107

**Authors:** Cynthia J. Downs, Laura A. Schoenle, Eric W. Goolsby, Samantha J. Oakey, Ray Ball, Rays H.Y. Jiang, Kirk C. Klasing, Lynn B. Martin

## Abstract

Terrestrial mammals span 7 orders of magnitude in body size, ranging from the < 2 g pygmy shrew (*Suncus etruscus*) to the > 3900 kg African elephant (*Loxodonta africana*). Although body size has profound effects on the behavior, physiology, ecology, and evolution of animals, how investment in immune defenses changes with body size is unknown. Here, we develop a novel 12-point dilution-curve approach to describe and compare antibacterial capacity against 3 bacterial species among >160 terrestrial species of mammals. We show that antibacterial activity in serum across mammals exhibits isometry, but the serum of large mammals is less hospitable to bacteria than would be predicted by their metabolic rates. Specifically, hypometric metabolic rates would predict that a large species should have disproportionately lower antibacterial capacity than small species, but body size is unrelated to killing capacity across species. Scaling of antibacterial immune defenses provides novel perspectives on the ecology of host-pathogen interactions, and on their co-evolutionary dynamics. These results have direct implications for effectively modeling the evolution of immune defenses and identifying potential reservoir hosts of zoonotic pathogens.

## Introduction

Hosts can be thought of as islands whose size impacts how many parasites, including microbes, can colonize them (Kuris et al. 1980). Bigger hosts represent bigger islands, leading parasite abundance (Kieft and Simmons 2015), biomass (Poulin and George-Nascimento 2007), diversity (Ezenwa 2004), size (Harrison 1915), size diversity (Poulin 2007), and prevalence (Ishtiaq et al. 2017) to increase with host body size. Large-bodied species also appear more susceptible to some parasite infections (Filion et al. 2020). Large body size is also intimately connected to the relative value of self-defense versus reproduction (Calder 1984), as most hosts reach large size via a long lifespan (Harrison 2017; Downs et al. 2020). Altogether, natural selection should favor the evolution of robust immune systems in large hosts (Sheldon and Verhulst 1996; Downs et al. 2014), and despite a few theoretical (Langman and Cohn 1987; Wiegel and Perelson 2004; Banerjee et al. 2017) and empirical (Lee 2006; Schoenle et al. 2018) calls to investigate scaling of immune defenses, very little works has yet considered whether and how immune defenses scale across the full range of terrestrial mammalian body sizes.

Generally, body size scaling relationships are modeled as power laws, *Y* = *a M^b^*, with linear transformations, log_10_ *Y* = log_10_ *a* + *b* log_10_ *M*, where *Y* is the focal trait, *M*is body mass, *a* is the intercept, and *b* is the scaling factor. Four hypotheses make predictions about the values of the scaling factors describing the relationship between immune defenses and body mass. To date, the Protecton and the Complexity of Immunity Hypotheses have received the most attention. Both hypotheses predict that immunity should scale in direct proportion to body size (i.e., isometrically), which results in a mass-invariant relationship (*b* = 0) for concentrations (Langman and Cohn 1987; Wiegel and Perelson 2004). The fractal nature of the circulatory and lymphatic systems is argued to predict isometry because rates of parasite detection and delivery of defenses should be driven by transit of cells and proteins through the vasculature (e.g., Wiegel and Perelson 2004; Banerjee et al. 2017; Banerjee 2018). Although this framework was developed for predicting the number of a single clone of B cells necessary to detect an antigen in a host and circulation time of a B cell (Wiegel and Perelson 2004), it has been extended to make predictions about other types of immune defenses but particularly adaptive immunity (Banerjee et al. 2017; Banerjee 2018). By contrast, the Rate of Metabolism Hypothesis predicts a hypometric coefficient for immunity (*b* < 0) because all cellular activities are linked to basal metabolic rates (BMR) (Brown et al. 2004). As mass-specific BMR scales at *b* = −0.25, immune defenses involving cell turnover and metabolism should scale similarly (Dingli and Pacheco 2006).

Finally, the Safety Factor Hypothesis invokes performance-safety relationships from biophysics (Harrison 2017) and physiology (Diamond 2002), and posits that large species should evolve disproportionately stronger (i.e., hypermetric, *b* > 0) damage-mitigation mechanisms than small ones (Harrison 2017; Downs et al. 2020). Immunologically, the Safety Factor Hypothesis predicts that large hosts should evolve exceptionally robust constitutive defenses to protect against the greater infection risks they will experience. Variants of the Safety Factor Hypothesis exist in the literature already, including Peto’s paradox for cancer (Peto 1977) and optimal defense theory (Shudo and Iwasa 2001). Only recently, however, has direct empirical evidence been provided for immune defenses, namely hypermetric scaling for both neutrophil concentrations in mammals (*b* = 0.11) (Downs et al. 2020) and heterophil concentrations in birds (*b* = 0.19) (Ruhs et al. 2020).

Despite evidence for some immune allometries, it remains obscure whether the functional capacity of immune defenses scale similarly. Does hypermetric scaling of granulocyte concentrations represent greater immune protection in large animals or simply evidence that large animals must compensate for lower per-granulocyte effectiveness by circulating more cells? Cell size scales isometrically with body mass, but per-gram metabolism scales hypometrically (Schmidt-Nielsen 1984), so the average large-animal leukocyte might be comparatively ineffective compared to a leukocyte from a small species. Moreover, as selection tends to act on integrated traits (Bennett and Huey 1990), scaling inference based on processes that affect host fitness (i.e., control of bacterial infections) will inherently be more insightful than studies solely based on cell concentrations. Few scaling studies of immunity to date have relied on integrated immune functions, and others either involved few species, narrow taxonomic groups, or found body mass effects but did not estimate allometries (e.g., Nunn 2002; Blount et al. 2003; Nunn et al. 2003; Lee et al. 2008; Schneeberger et al. 2013; Tian et al. 2015; Fassbinder-Orth et al. 2019; reviewed in Downs et al. 2020).

Our goals here were i) to determine how blood-based antibacterial defenses scaled with body size among terrestrial mammals, and ii) to test which hypothesis best-predicts the scaling exponent (*b*) for this type of immunity. We focused on antibacterial capacity of sera (French and Neuman-Lee 2012), as this defense is functional and integrative (Demas et al. 2011), is an important determinant of host health and virulence for certain bacterial pathogens (Taylor 1983), and is measurable in a standard, reagent-independent way. Briefly, antibacterial capacity measures bacteriostasis and bacteria killing capacity of sera. To describe interspecific variation in this immune function in the most confound-free manner possible, we compared the efficacy of sera of several, independent replicates from adults of mammalian species housed in zoos. To provide generality to any scaling we observed, we quantified the antibacterial capacity of sera of 160+ mammalian species against three distinct bacteria, *Escherichia coli* (ATCC #8739),

*Salmonella enterica* (ATCC #13311), and *Micrococcus luteus* (ATCC # 4698). These bacteria are common pathogens of many vertebrates (Mastroeni et al. 2001), some portion of their natural infection timeline occurs in blood (Mittrücker and Kaufmann 2000), and they are controlled effectively by blood-borne immune defenses (Taylor 1983). As the last common evolutionary ancestor of our focal microbes was ancient, any consistency in allometries should be more attributable to host immunity than parasite evolutionary history.

To enable meaningful comparisons across species, we characterized antibacterial dilution series, instead of antibacterial capacity at single dilutions (as is typically used, French and Neuman-Lee 2012), as the shapes of resultant *ex vivo* antibacterial functions capture salient aspects of defense (Fig. 1A). For instance, if a very dilute serum sample was effective against a standard amount of bacteria, a given individual of that host species would be relatively adept at controlling a large initial dose of bacteria in its bloodstream. Other meaningful antibacterial parameters included i) the maximum amount of bacteria that serum could protect against, ii) the serum dilution at the point of most rapid change from protected to vulnerable, and iii) the slope at this steepest part of the antibacterial function (Fig. 1A). This last parameter could be a proxy for either how rapidly serum defenses could be recruited to a bacterial replication event (Murphy et al. 2007) or capture the pattern by which proteins and other molecules were recruited combinatorically to control bacteria. The Safety Factor Hypothesis, our focal hypothesis, predicts that large mammals will have disproportionately higher maximal antibacterial capacity, higher antibacterial capacity at a specific dilution, and a more rapid shift from vulnerable to complete protection than small species (Fig. 1B). By contrast, the Rate of Metabolism Hypothesis predicts the opposite for large mammals relative to small ones (Fig 1C), and the Protecton and Complexity Hypotheses predict that large mammals will have the same antibacterial capacity as small species (Fig. 1D).

**Figure 1.**
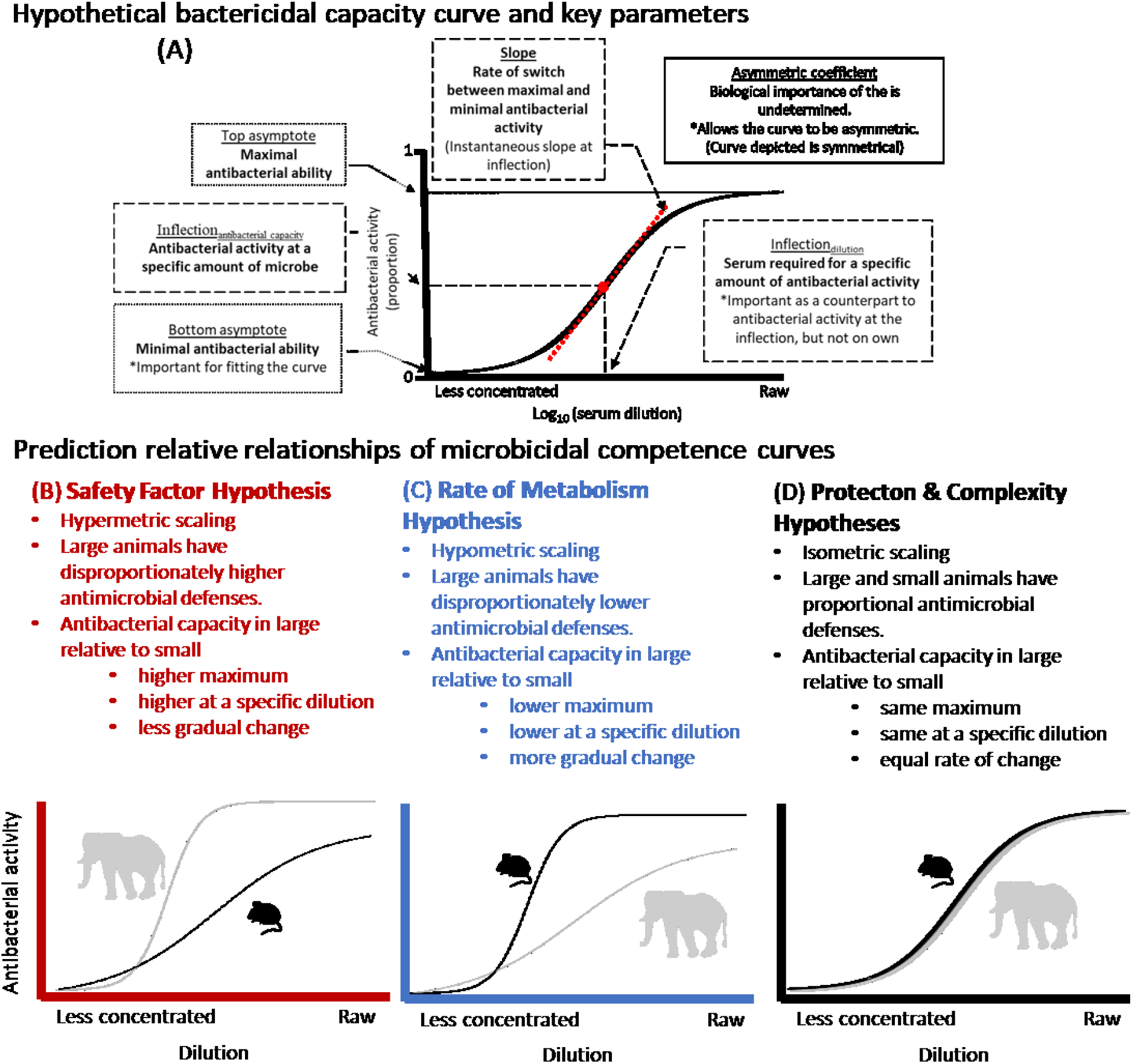
Antibacterial capacity dilution curves support the Safety Factor Hypothesis. Hypothetical antibacterial dilution curve showing the parameters we estimated and biological interpretation (A) This curve shows, names (underlined), and described the curve parameters and the biological interpretation of the curve parameter (in bold). Hypothetical antibacterial curves for an elephant and mouse showing predictions for the Safety Factory Hypothesis (B), the Rate of Metabolism Hypothesis (C), and the Protecton and Complexity Hypotheses (D).

## Materials and Methods

### Samples

We used the simulations performed by Dingemanse and Dochtermann (Fig. 1 of 2013) to guide our sample sizes. We obtained serum from healthy, zoo- and lab-housed animals ranging in body size from 16 g – 3,600 kg, which covers the range of body sizes of extant terrestrial mammals (Fig. S1) (Smith and Lyons 2011). We supplemented these samples, which comprised the vast majority of our sample, with those from lab-housed animals to augment the small-bodied species (< 2 kg) included in our analysis, as these body sizes are rare for mammals in most zoos. Most zoo samples were collected as part of routine wellness checkups whereas lab samples were collected at the termination of experiments; all animals from which samples were taken were outwardly healthy. We could not run all assays against all bacterial species for all samples because of sample volume constraints. Final sample sizes of mammal species were 199, 186, and 164 for antibacterial capacity against *E. coli*, *S. enterica*, and *M. luteus*, respectively. We assayed a mean of ≥ 4 samples per species for each microbe (n = 1-41 replicates per species, Fig. S2). We also obtained >15 samples from a subset of species representing the full range of body masses to ensure that a single individual from a large or small species was not biasing a species mean (Fig. S3). Samples were stored at the collection location at −80 to −20 °C until shipped on dry ice to our research labs where they were stored at 80 °C until assays. Samples were used in assays within 24h of thawing. We restricted our analysis to samples collected in the years 2005-2019 because preliminary work indicated that older samples had reduced antibacterial ability. Use of these samples was approved by the Institutional Animal Care and Use Committees at home institutions. Samples were collected as part of routine veterinary exams under auspices of zoo IACUCs and were shared after approval by the appropriate committee at each zoo.

### Antibacterial capacity

We measured antibacterial capacity against *E. coli* (ATCC #8739), *S. enterica* (ATCC #13311), and *M. luteus* (ATCC # 4698) using an adaptation of the microbiocidal assay (a.k.a. bacteria killing assay, BKA, microbiocidal activity) developed by French and Neuman-Lee (2012). Briefly, we plated a 12-point dilution curve of each serum sample in triplicate on a round-bottomed 96-well plate. *E. coli* had two serial dilutions: 6 points from undiluted serum to 1:64, and 6 points from 3:4 to 3:256. *S. enterica* also had two serial dilutions: 6 points from undiluted to 1:64, and 6 points from 3:4 to 3:128. *M. luteus* had a single serial dilution ranging from undiluted serum to 1:2048. Super antibacterial capacity curves—curves for species with 100% antibacterial capacity at the lowest dilution of the normal curves—for *E. coli* were 6 points from 1:4 to 1:256 and 6 points from 3:16 to 3:512 and for were *S. enterica:* 6 points from 1:16 to 1:512 and 6 points from 3:4 to 3:128. We also plated a 4-point dilution curve in Dulbecco’s Phosphate Buffered Saline (PBS, Sigma-Aldrich #D8537) of commercially available cow serum (Innovative Research Novi, MI 48377, #IBV-Compl) as our inter-assay control. Cow curve dilutions were 1:32, 1:64, 1:128, 1:256 for *E. coli*; 1:16, 1:32, 1:64, 1:128 for *S. enterica;* and 1;20, 1:80, 1:320, 1:1280 for *M. luteus*. Our final volume of each serum dilution was 18 μl. We plated three replicates of a 20 μl negative control of PBS and 9 replicates of an 18 μl positive control. We added 2 μl of a standard concentration (10^4^ CFU ml^-1^) of bacteria to all wells except the negative controls. We incubated plates for 30 min at 37 °C. We next added 125 μl of tryptic soy broth (BD #211825) to all wells, and then shook plates for 1 min at 300 rpm. We then measured baseline absorbance of all wells at 300 nm (Biotek Synergy HTX Multi-Mode Reader) to serve as an internal control before incubating covered plates for 12 h for *E. coli*, 10 h for *S. enterica*, and 48 h for *M. luteus* at an incubation temperature of 37 °C. We then measured final absorbance of all wells again at 300 nm and calculated antibacterial capacity for each well as 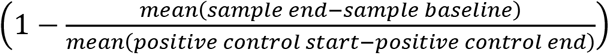 * 100%. Inter-assay (9.02%) and intra-assay variation (0.02%) were both low. Key parameters of the antibacterial capacity assay for each microbe are summarized in Table S1 and full details about the procedures used to develop and validate our assays are available in the Supplementary Methods. Assay protocols are deposited on FigShare (Schoenle et al. 2020).

### Data processing and analysis

All data processing and analyses were performed in the statistical programming language R (R Core Team 2019). We compiled data using a workflow developed using R (all code files are available on Figshare (Keith et al. 2020; Downs et al. 2021)). We visually checked each sample dilution curve for outliers (i.e., replicates that were distinct from the other two replicates and/or fell off the expected sigmoidal curve). We determined whether positive controls, negative controls, and inter-assay controls (i.e., cow sera) were within the expected range, and discarded a plate if data fell outside expected ranges and reran samples.

To identify any allometries, we first fit non-linear antibacterial functions for each individual sample separately. Properly estimating allometries required models that allowed us to perform phylogenetically multivariate regressions while accounting for the sampling of numerous individuals within a species because we know that species are not independent samples (Felsenstein 1985) and a failure to account for within-species variation can produce inaccurate regression estimates (Garamszegi and Møller 2010; Garamszegi 2014). This modeling framework is under-served by the available statistical packages, so we modeled data using two main approaches. For our first approach, we used multivariate phylogenetic comparative analyses of our data using the R package `Rphylopars` (Goolsby et al. 2021). This approach allowed us to calculate phylogenetic covariances of traits among species while accounting for numerous observations within a species (i.e., among-individual variation) (Goolsby et al. 2017). Although this approach did not estimate scaling coefficients as they would be estimated with a regression/linear model-based approach, it allowed us to estimate phylogenetic signal of antibacterial capacity and correlations among the antibacterial curve parameters and between curve parameters and body mass in the same model. For our second approach, we built multivariate general linear mixed models using the R package `MCMCglmm` (de Villemereuil and Nakagawa 2014). Although these models assume each species is an independent point, they allowed us to estimate *b* for all curve parameters simultaneously while accounting for the variance within species by including multiple observations per species variation (Dingemanse and Dochtermann 2013). This approach better parallels the traditional approach to estimating scaling coefficients using regression models (Sieg et al. 2009). To test for the strength of phylogenetic signals in a framework based on general linear models, we built phylogenetic univariate general linear mixed models for each antibacterial curve parameter individually using `MCMCglmm` (supplementary methods) (de Villemereuil and Nakagawa 2014). We extracted species relatedness from a tree build using PhyloT (Letunic 2015, Fig S4)

#### Fitting non-linear curves for each individual

For the first step, we fit 5-parameter logistic regression growth curves to each individual antibacterial curve (i.e., the shape of dilution curves across all 12-serum concentrations) using package `nplr` (Commo and Bot 2016) (Fig. 1A). To aid in curve-fitting, we log10-transformed serum concentrations (i.e., the dilutions) and converted the antibacterial ability from a percent to a proportion. Curves could only be fit to values between 0 and 1. So percent inhibited values > 100 were forced to a random value between 99 and 100 and percent inhibited values < 0 were forced to a random value between 0 and 1 before conversion to proportions. This approach restricted values to a range near maximal and minimal antibacterial activity for all other curves. We extracted the curve parameters (inflection_dilution_, inflection_antibacterial capacity_, bottom asymptote, top asymptote, slope, asymptote coefficient) to use in univariate and multivariate general linear models (Fig. 1A). Slope and inflection_dilution_ were already on a log_10_-scale. We added 1 to inflection_antibacterial capacity_, bottom asymptote, top asymptote and then log_10_-transfromed them. We also log_10_-transformed body mass. All analyses were performed using transformed data.

#### Phylogenetic covariance

We performed a multivariate phylogenetic comparative analysis for antibacterial capacity against each microbe using the R package `Rphylopars` (Goolsby et al. 2021). All antibacterial capacity curve parameters and body mass were included as response variables in these models, with one exception. Top asymptote was not included in the model for antibacterial capacity against *S. enterica* because it was highly correlated (r = 0.998) with Inflection_antibacterial capacity_; both could not be included in the same model. We included phylogenetic effects from a tree we produced by pruning the time-rooted phylogenetic tree created by Uyeda et al. (2017) to our species list; we scaled tree with height scaled to 1. Models were fit assuming a Brownian motion model of evolution and we estimated the percent of variance explained by phylogeny. We tested for phylogenetic signal for each trait using a likelihood ratio test (df = 1) to compare Pagel’s lambda against a null model with a star phylogeny (lambda = 0) to our model that included phylogeny (Pagel 1997, 1999).

#### Multivariate mixed models

We generated separate multivariate mixed models to query body mass effects on antibacterial capacity, one for each bacterial species. All curve parameters were included as response variables, and we allowed each parameter to have different slopes and intercepts. Body mass was included as a fixed effect, and species was incorporated as a random effect. Species had independent intercepts and slopes. These models were fit using the `MCMCglmm` package (Hadfield 2010; Hadfield and Nakagawa 2010; de Villemereuil and Nakagawa 2014). All mixed models were fit using a weak inverse-Gamma prior with shape and scale parameters set to 1.002 for the random effect. Default priors for all other fixed effects were used. Model chains were run for 1.82 × 10^6^ iterations, a 420,000-iteration burn-in, and a 1400 iteration thinning interval. Results were robust across alternative priors, and chain length was sufficient to yield negligible autocorrelation. We extracted the slopes describing the relationship between each parameter and body mass across all species.

We included bacterial capacity at the bottom asymptote and asymmetric coefficient in our models because they helped describe antibacterial capacity curves, although we did not make predictions about their biological functions. In our study, we did not always collect data at the lower asymptote because we limited our sampling to 12-dilutions. As a result, the bottom asymptote estimated by the general linear model was based on incomplete data and not biologically relevant to our scaling hypotheses. The dilution at the inflection point was also included in our models because it informed our understanding of the antibacterial capacity at the inflection point, but it was not obviously biologically meaningful for understanding immune scaling.

## Results

Overall results from all three modeling approaches were consistent and supported isometric (i.e., mass-invariant) scaling (*b*=0) for almost all curve parameters for antibacterial capacity parameters against all three microbes. The one exception to this pattern was the slope for antibacterial capacity against *M. luteus*; in this one instance, all three modeling approaches indicated a negative relationship between slope at the intercept with body mass such that slopes become less steeply positive with body mass.

### Phylogenetic covariance

Models built using `Rphylopars` estimated that the covariance between body mass and any of the curve parameters was very close to zero and had 95% CI intervals that overlapped zero, with one exception (Table 1): the correlation between slope and body mass for antibacterial capacity against *M. leuteus* had a negative correlation of −11.0 (95% CI: −22.8: −2.6). Lambda values provided evidence of phylogenetic signal in body mass and at least one curve parameter for antibacterial capacity against all three microbes (Table 2). Phylogeny explained >97% for variation in body mass, but only ≤ 30% of the variation in most of the antibacterial capacity curve parameters (Table 2). An exception was that phylogeny explained 54% if of the variation in Inflection_dilution_ for antibacterial capacity against *S. enterica*. Phylogenetic correlations among curve parameters and phylotypic correlations can be found in Tables S1 and S2, respectively. These models had slightly smaller sample sizes because not all species were in the time-rooted tree. Final sample sizes of species for these models were 191, 178, and 157 for antibacterial capacity against *E. coli, S. enterica*, and *M. luteus*, respectively

**Table 1.** Covariance (Mean ± 95% CI) between body mass and antibacterial capacity curve parameters from phylogenetic covariance models estimated using the `Rphylopars` in Program R.

| Antibacterial capacity curve parameter | Estimate of covariance by dataset |  |  |
| --- | --- | --- | --- |
|  | *E. coli* | *S. enterica* | *M. luteus* |
| Top asymptote<sup>#</sup> | -0.002<br>(-0.036 : 0.035) | N/A* | -0.024<br>(-0.113 : 0.064) |
| Slope | 9.50<br>(-4.3 : 26.3) | -0.394<br>(-11.9 : 12.6) | -11.0<br>(-22.8 : -2.6) |
| Inflection<sub>dilution</sub> | -0.262<br>(-0.723 : 0.360) | -0.126<br>(-0.747 : 0.465) | -0.103<br>(-0.954 : 0.947) |
| Asymptote coefficient | -0.022<br>(-0.345 : 0.178) | -0.107<br>(-0.341 : 0.033) | -0.022<br>(-0.390 : 0.156) |
| Inflection<sub>antibacterial capacity</sub> | -0.006<br>(-0.284 : 0.036) | 0.009<br>(-0.146 : 0.445) | -0.023<br>(-0.087 : 4.47) |
| Bottom asymptote | -0.002<br>($-1.3 \times 10^{78}$ : $2.1 \times 10^{95}$) | -0.009<br>(-0.409 : 0.046) | -0.041<br>($-2.0 \times 10^{33}$ : 3.61) |

<sup>#</sup> Inflection<sub>antibacterial capacity</sub>, top asymptote, and bottom asymptote were +1, log<sub>10</sub>-transformed. Body mass and slope were log<sub>10</sub>-transformed prior to beta estimation.
\* Top asymptote was not included in the model for antibacterial capacity against *S. enterica* because it was highly correlated ( $r = 0.998$ ) with Inflection<sub>antibacterial capacity</sub>. Both variables could not be included in the same model

**Table 2.**
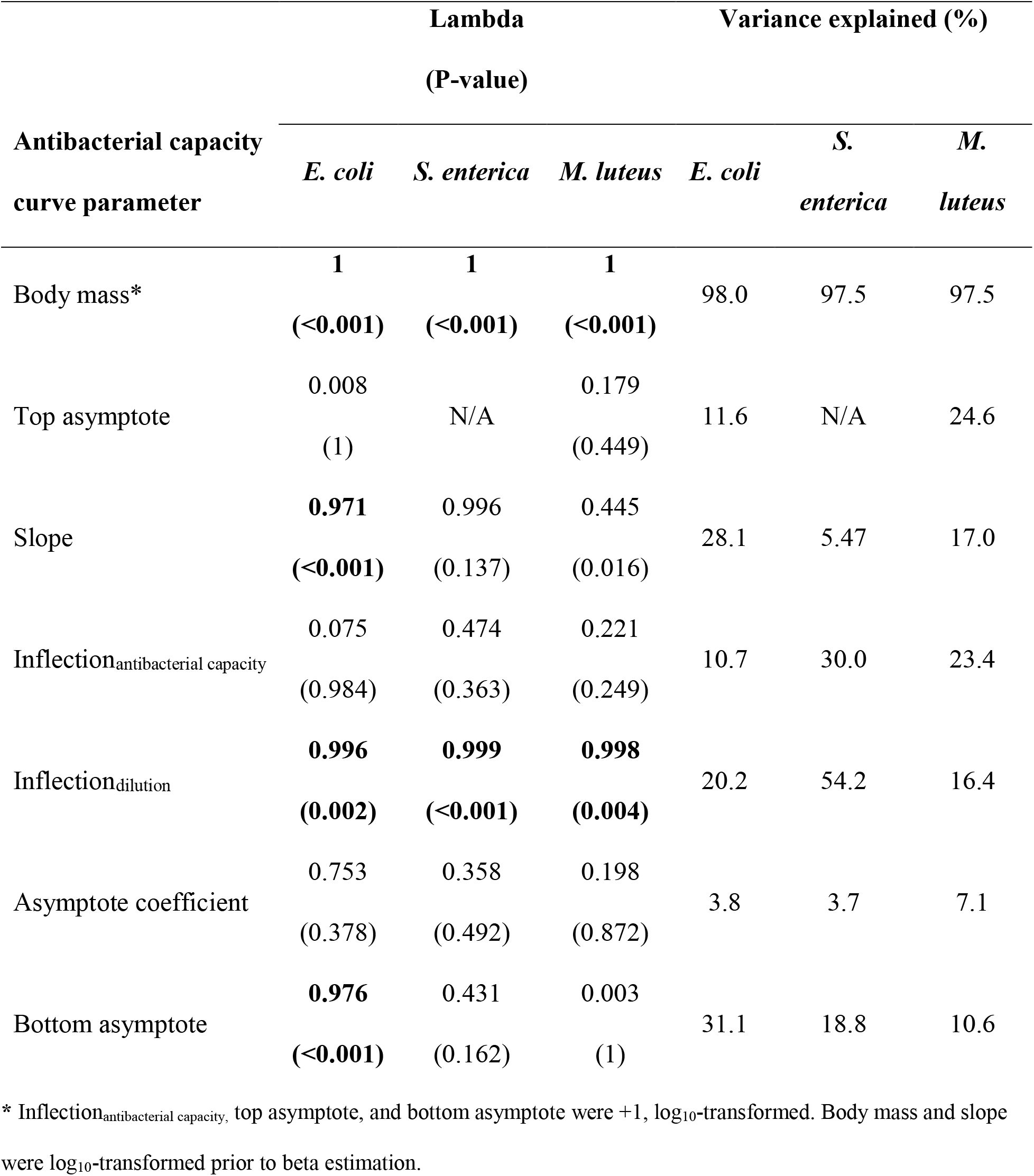
Estimates of the percent of variation in antibacterial curves against *E. coil, S. enterica*, and *M. luteus* explained by phylogeny from phylogenetic covariance models estimated using the `Rphylopars` in Program R.

### Phylogenetic univariate mixed effects models

Phylogenetic univariate models supported scaling coefficients of zero. The 95% CI for all curve parameters for antibacterial capacity against *E. coli*, *S. enterica*, *M. luteus* overlapped zero except for slope of antibacterial curves against *M. luteus* (supplemental materials, Table S3). This parameter scaled hypometerically (−4.30, 95% CI: −8.34, −0.30). Phylogeny explained 1-18% of the variation in antibacterial curve parameters (supplemental materials, Table S4).

### Multivariate mixed effects models

Multivariate models supported scaling coefficients of zero for all curve parameters for antibacterial capacity against *E. coli*, *S. enterica*, *M. luteus* with two exceptions (Table 3, Fig. 2) The slope of antibacterial curves against *M. luteus* had a hypermetric slope of −4.3 (95% CI: −7.7: −0.79) and the asymmetric coefficient scaled with a slope of −3.78 (95% CI: −5.604: −0.91).

**Figure 2.**
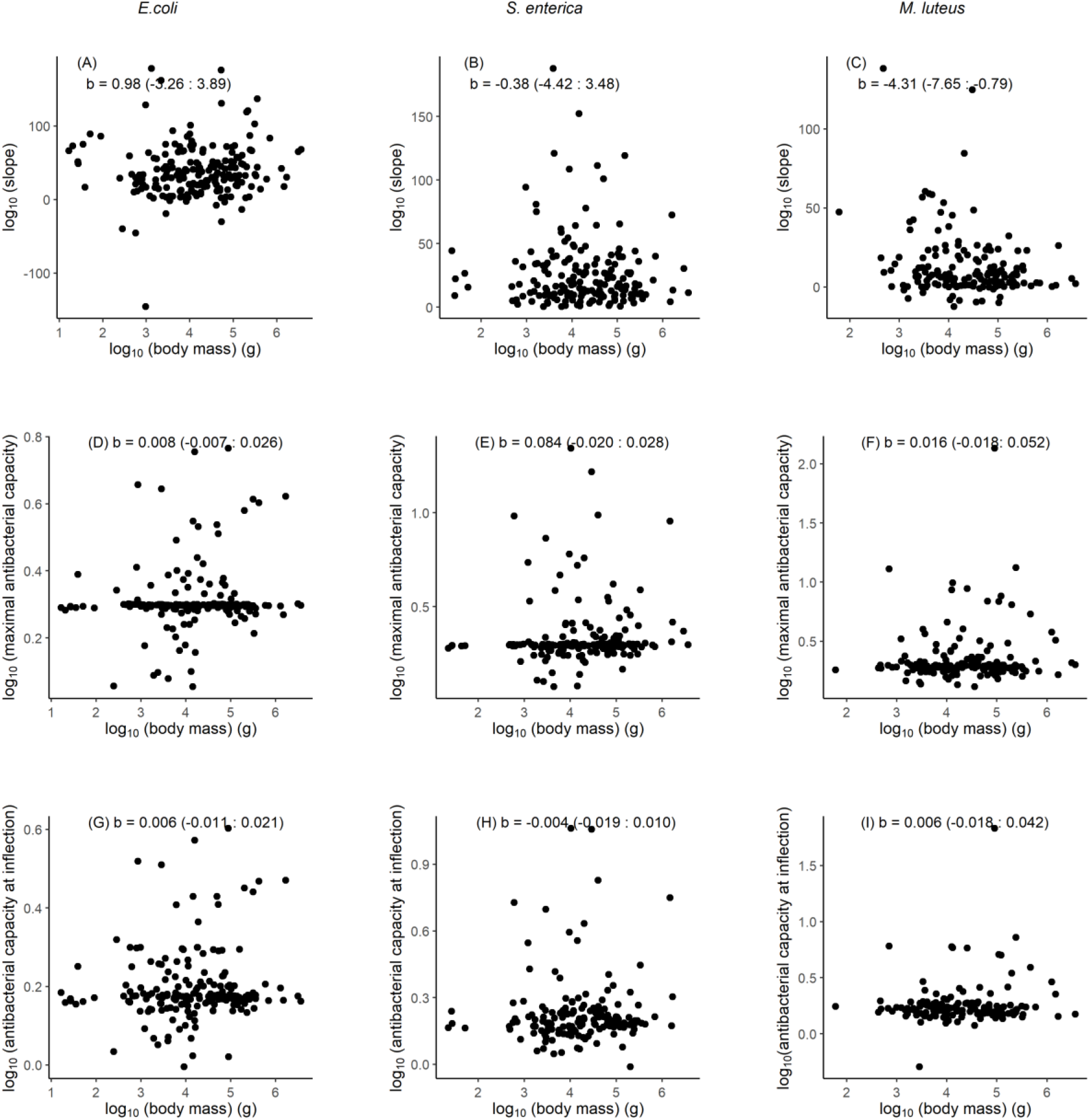
Parameters of antibacterial capacity curves—instantaneous slope at the inflection point (A-C) maximal antibiotic capacity (D-F), and antibiotic capacity at the inflection point (G-I)—against *Escherichia coli* (A, D, G,), *Salmonella enterica* (B, E, H), *Micrococcus luteus* and (C, F, I) against body mass. Each point represents a species mean, but analyses were performed on individual-level data.

**Table 3.**
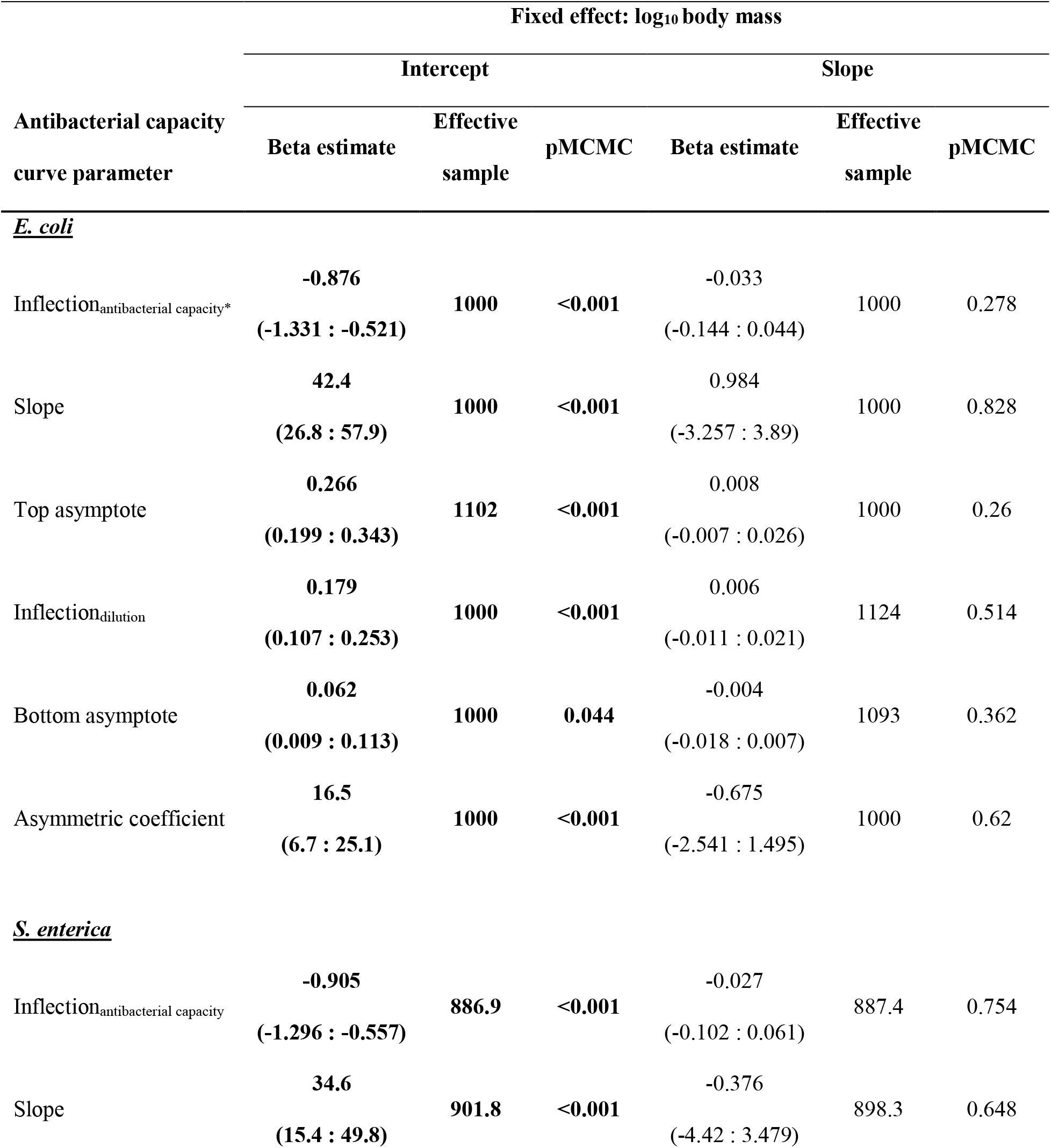
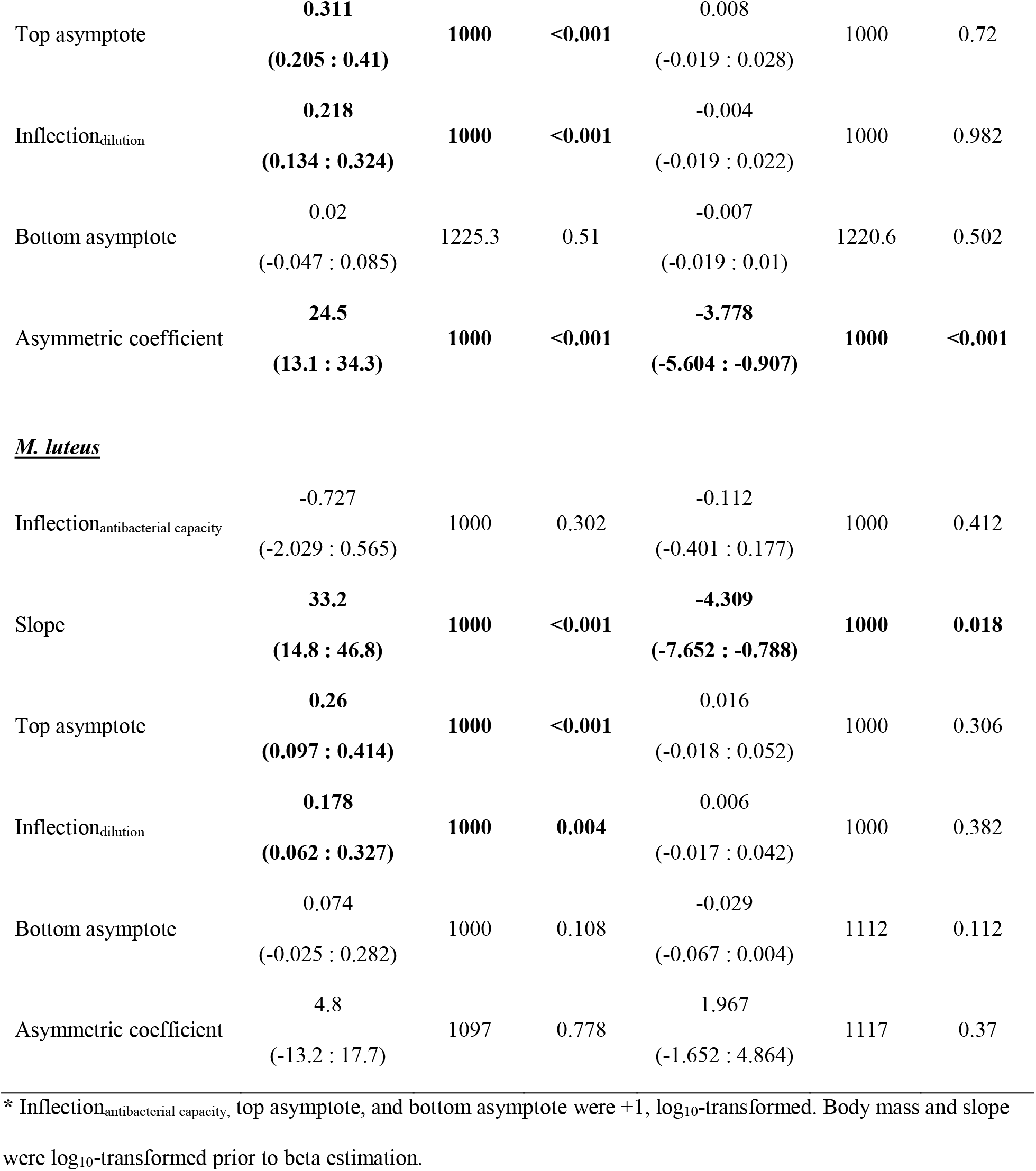
Estimated intercepts and scaling coefficients (mode ± 95% CI) for curve parameters of antibacterial capacity against *E. coli*, *S. enterica*, *M. luteus* from phylogenetic univariate mixed effects models. Estimates in **bold** have 95% CI intervals that do not overlap 0.

| Antibacterial capacity<br>curve parameter | Fixed effect: log<sub>10</sub> body mass |  |  |  |  |  |
| --- | --- | --- | --- | --- | --- | --- |
|  | Intercept |  |  | Slope |  |  |
|  | Beta estimate | Effective sample | pMCMC | Beta estimate | Effective sample | pMCMC |
| <u>*E. coli*</u> |  |  |  |  |  |  |
| Inflection<sub>antibacterial capacity*</sub> | **-0.876**<br>(-1.331 : -0.521) | **1000** | **<0.001** | -0.033<br>(-0.144 : 0.044) | 1000 | 0.278 |
| Slope | **42.4**<br>(26.8 : 57.9) | **1000** | **<0.001** | 0.984<br>(-3.257 : 3.89) | 1000 | 0.828 |
| Top asymptote | **0.266**<br>(0.199 : 0.343) | **1102** | **<0.001** | 0.008<br>(-0.007 : 0.026) | 1000 | 0.26 |
| Inflection<sub>dilution</sub> | **0.179**<br>(0.107 : 0.253) | **1000** | **<0.001** | 0.006<br>(-0.011 : 0.021) | 1124 | 0.514 |
| Bottom asymptote | **0.062**<br>(0.009 : 0.113) | **1000** | **0.044** | -0.004<br>(-0.018 : 0.007) | 1093 | 0.362 |
| Asymmetric coefficient | **16.5**<br>(6.7 : 25.1) | **1000** | **<0.001** | -0.675<br>(-2.541 : 1.495) | 1000 | 0.62 |
| <u>*S. enterica*</u> |  |  |  |  |  |  |
| Inflection<sub>antibacterial capacity</sub> | **-0.905**<br>(-1.296 : -0.557) | **886.9** | **<0.001** | -0.027<br>(-0.102 : 0.061) | 887.4 | 0.754 |
| Slope | **34.6**<br>(15.4 : 49.8) | **901.8** | **<0.001** | -0.376<br>(-4.42 : 3.479) | 898.3 | 0.648 |
| Top asymptote | **0.311**<br>**(0.205 : 0.41)** | **1000** | **<0.001** | 0.008<br>(-0.019 : 0.028) | 1000 | 0.72 |
| Inflection<sub>dilution</sub> | **0.218**<br>**(0.134 : 0.324)** | **1000** | **<0.001** | -0.004<br>(-0.019 : 0.022) | 1000 | 0.982 |
| Bottom asymptote | 0.02<br>(-0.047 : 0.085) | 1225.3 | 0.51 | -0.007<br>(-0.019 : 0.01) | 1220.6 | 0.502 |
| Asymmetric coefficient | **24.5**<br>**(13.1 : 34.3)** | **1000** | **<0.001** | **-3.778**<br>**(-5.604 : -0.907)** | **1000** | **<0.001** |
| **M. luteus** |  |  |  |  |  |  |
| Inflection<sub>antibacterial capacity</sub> | -0.727<br>(-2.029 : 0.565) | 1000 | 0.302 | -0.112<br>(-0.401 : 0.177) | 1000 | 0.412 |
| Slope | **33.2**<br>**(14.8 : 46.8)** | **1000** | **<0.001** | **-4.309**<br>**(-7.652 : -0.788)** | **1000** | **0.018** |
| Top asymptote | **0.26**<br>**(0.097 : 0.414)** | **1000** | **<0.001** | 0.016<br>(-0.018 : 0.052) | 1000 | 0.306 |
| Inflection<sub>dilution</sub> | **0.178**<br>**(0.062 : 0.327)** | **1000** | **0.004** | 0.006<br>(-0.017 : 0.042) | 1000 | 0.382 |
| Bottom asymptote | 0.074<br>(-0.025 : 0.282) | 1000 | 0.108 | -0.029<br>(-0.067 : 0.004) | 1112 | 0.112 |
| Asymmetric coefficient | 4.8<br>(-13.2 : 17.7) | 1097 | 0.778 | 1.967<br>(-1.652 : 4.864) | 1117 | 0.37 |

\* Inflection<sub>antibacterial capacity</sub>, top asymptote, and bottom asymptote were +1, log<sub>10</sub>-transformed. Body mass and slope were log<sub>10</sub>-transformed prior to beta estimation.

## Discussion

Our data provide consistent and robust evidence for isometric (i.e., mass-invariant) scaling of serum antibacterial activity in terrestrial mammals. Additionally, results indicate that phylogenic signal was surprisingly low for all three microbes. Superficially, isometric scaling supports the prediction of the Protection and Complexity Hypotheses. These results were somewhat surprising, given that large species face a greater risk of infection and face greater lost fitness costs from infections (Downs et al. 2020). Below we argue that the interpretation of these results might support a more complex interpretation once integrated with our understanding of how host metabolic rates and other immunological defenses scale.

Mass-invariance of antibacterial capacity was consistent regardless of the bacterial species studied. This discovery is interesting in light of the ecology and life history differences among bacterial species. *M. luteus* grows slowly and causes modest disease in most hosts relative to the other two bacteria; both *S. enterica* and *E. coli* can cause sepsis (van der Poll and Opal 2008), infect via the diet, and have similar replication rates in culture. Our findings thus support the idea that broadly effective, fast-acting immune responses are generally favored, rather than by a specific pathogenetic target, across mammalian species.

### Isometric scaling and the Protecton/Complexity Hypotheses

Both of the Protecton and Complexity Hypotheses hypotheses argue that large and small animals need equal defenses against parasites, resulting in immune responses having a scaling exponent of 0 (Langman and Cohn 1987; Wiegel and Perelson 2004; Banerjee and Moses 2010). Our results are consistant with this prediction. However, by definition, isometric scaling is difficult to test because it corresponds with a null model of a mass-invariant or intercept-only model. Statistically, we cannot distinguish between the null and the model prediction.

Perhaps more importantly, both the Protecton and Complexity Hypotheses were developed to predict how the number of lymphocyte clones should scale with body size to provide protection against foreign antigens (Langman and Cohn 1987; Wiegel and Perelson 2004; Banerjee and Moses 2010), not to explain constitutive, innate immunity as studied here. Assumptions about the distribution and timing of adaptive immune responses are unlikely to hold for the early-acting, constitutive innate immune responses. The adaptive and innate immune systems operate on different time scales (Murphy et al. 2007), and the life cycles of lymphocytes and neutrophils are quite different. Circulating peripheral neutrophils have a lifespan of 1-7 days (Hidalgo et al. 2018) whereas peripheral, naïve lymphocytes are long lived—T cells live on the order of years and B cells have a half-life of 4 weeks to several months—and later phases of adaptive immunity (i.e., memory) are controlled by even longer-lived lymphocytes (Tough and Sprent 1995). Even “short-lived” lymphocytes involved in early stages of memory have life-spans of weeks, which is much longer than neutrophils (Tough and Sprent 1995). Although the Complexity Hypothesis has been extended to make predictions about the size and number of lymph nodes in mammals (Banerjee 2018) and host capacity to replicate West Nile virus (Banerjee and Moses 2009, 2010; Banerjee et al. 2017), these extensions are still primarily based on the dynamics of adaptive immunity. It follows that the underpinning logic of the Complexity Hypothesis would need to be rederived from first principles to account for dynamics of innate immune defense and to make predictions for early, broadly active defenses that are part of the innate immune system, such as antibacterial capacity.

### Isometric scaling is steeper than expected by metabolic rates

The mass-invariance of antibacterial capacity is intriguing when considered with the well-documented hypometric scaling of mass-specific metabolic rates (Brown et al. 2004; Savage et al. 2004) and the hypermetric scaling of concentrations of some immune cell types (Downs et al. 2020). The Metabolic Theory of Ecology posits that mass-specific metabolic rate can be interpreted as the ratio of the average metabolic rate of a cell to the average cell volume, and this ratio also has a scaling exponent of −0.25 (Savage et al. 2007):

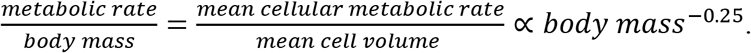

If cellular metabolic rate increases, cell volume must decrease (Savage et al. 2007). Thus, for cell types with mass-invariant volumes (b = 0), cellular metabolic rate will scale with the same slope as mass-specific metabolic rate (b = −0.25). It follows that if basal metabolic rate is the pacemaker for cellular activity, protein synthesis for these cell types would also scale with a slope of −0.25 (Brown et al. 2004).

Our assay measured the functional capacity of complement and other circulating proteins (i.e., lysozyme, mannan-binding protein, beta-defensins) to opsonize and lyse bacteria (Demas et al. 2011; French and Neuman-Lee 2012). Complement proteins are predominantly produced by hepatocytes (Lubbers et al. 2017), and hepatocyte volume is mass-invariant (Savage et al. 2007). As such, production of complement protein production per hepatocyte should scale at −0.25. Liver mass scales with a slope of 0.895 (Prothero 2015), so. if each hepatocyte produces complement proteins at the same rate, antibacterial capacity should scale at 0.645 (= 0.895-0.25). Within this framework, a scaling coefficient of 0 for antibacterial capacity, as we described, is shockingly high. In other words, large mammals have much higher antibacterial capacity than expected by their metabolic rate alone. An outstanding question is how large mammalian species obtain the same antibacterial capacity as small ones if they are constrained by their metabolic rates.

### Why isometric scaling?

Antibacterial capacity and neutrophil concentrations, the other type of first, constitutive lines of host defense thus far studied in an allometric context (b = 0.11; Downs et al. 2020), both scaling more steeply than expected by scaling of metabolic rates. These scaling repationships may have evolved because, relative to small mammals, large mammals (1) have a disproportionately higher exposure to parasites and (2) have a disproportionately larger gap in replication rates relative to invading microbes (Downs et al. 2020). We will walk through each in turn.

First, large mammals have different exposure risks to parasites because of their ecology. Relative to small mammals, large mammals consume more food (Nagy 2001; Nunn et al. 2003), move farther and thus are exposed to more of habitat with each unit of movement (e.g., step, wing flap) (Schmidt-Nielsen 1984), and they have larger home ranges which exposes them to more types of habitats (Lindstedt et al. 1986; Kelt and Van Vuren 2001). Although a given gram of tissue in a large host is less exposed to parasites because of a low ratio of surface area to volume, absolute exposure to parasites is probably much higher in large than small species (Dobson and Hudson 1986; Downs et al. 2019). Large mammals also have a higher absolute exposure to parasites over their lifetime because they have longer lifespans than small mammals (Peters 1983; Calder 1984).

Second, parasites have an evolutionary advantage over hosts because of differences in replication rates. When parasites grow rapidly, as bacterial (and viral) infections often do, defenses with short time-delays are favored (Shudo and Iwasa 2001). Barriers against all-comers are the best option when adversaries can replicate faster than defenses can be honed and mobilized. The downside to these forms of defense is that they impose comparatively high costs, especially when adversaries are absent (Sears et al. 2006). Recently, a meta-analysis revealed that small animal species that are relatively long-lived incur the highest costs of *induced* immune defenses (Brace et al. 2017). Given the tendency of small species to breed prolifically and develop rapidly, isometric allometries might manifest as much because small species favor rapidly-mobilizable defenses that are activated only after exposure to a parasite as they do because of mandates of large size.

Our understanding of how activation of adaptive immune responses occurs would also support the need for large animals to have higher early constitutive immune defenses than expected by metabolic rates. B and T cell receptor diversity is more constrained by over-responsiveness to self-than under-responsiveness to non-self (De Boer and Perelson 1993). Indeed, the probability that a parasite will evade the antigen space covered by B and T cell repertoires is 4×10^-44^ (Perelson and Oster 1979; Banerjee et al. 2017). Adaptive immune systems will, therefore, eventually detect any antigen that ever evolves. The challenge of relying solely on these defenses, though, is that they involve major time lags (i.e., minimally 4-10 days but perhaps longer if the immune system is still developing (Ricklefs 1992). Lags for inducible defenses generally leave hosts vulnerable as microbial doubling time proceeds over a much shorter period. This incompatibility might explain why among 25 avian species, the production rate of infectious virions and the burst size of cells (i.e., the number of infectious virions released by an infected cell over its infected lifespan) scaled hypometrically, yet the rate of antibody-mediated neutralization of virus was isometric (Banerjee et al. 2017). Perhaps isometrically-scaled constitutive immune defenses produced the surprisingly low viral outcomes observed in that study.

If considered more broadly, our results might in fact be consistent with the Safety Factory Hypothesis. Perhaps instead of predicting hypermetric scaling of all forms of constitutive innate immunity, the Safety Factory Hypothesis should instead posit that large and small animals need equal *functional* protection against pathogens resulting in hypermetric scaling of some defenses and isometric (or even hypometric) scaling of others. Large mammals (Downs et al. 2020) and large birds (Ruhs et al. 2020) have disproportionally higher concentrations of early responding granulocytes relative to their small counterparts. However, this hypermetric scaling may not translate into hypermetric scaling of functionally because the synthesis of immunological molecules and cellular rates of reaction might be constrained by metabolic rate (Brown et al. 2004; Savage et al. 2007). In other words, the disproportionately high concentrations of granulocytes in large species might be necessary to have similar functionality as small species (Downs et al. 2020; Ruhs et al. 2020). Large animals may use a strategy of many cells with reduced activity, whereas small animals may use a strategy of fewer cells with higher activity. To test further test this idea, we would need to determine the scaling coefficient for the functionality of neutrophils and the scaling of the number of hepatocytes that produce complement and their functionality.

### Phylogenetic signal and other influences on antibacterial capacity

Our results also demonstrated that mammals exhibit a high diversity of antibacterial curves (Movie S1) and that little of this variation is explained by phylogeny or body size. The weak phylogenetic signal and a small percentage of variation in antibacterial curves explained by phylogeny in general were surprising. Previous analyses of scaling of leucocyte concentrations and in mammals (Downs et al. 2020) and birds (Ruhs et al. 2020) and leucocyte ratios in birds (Minias 2019) demonstrated that phylogeny explained the majority of interspecific variation. Although phylogenetic signal was weak for many aspects of antibacterial capacity we measured, we caution against equating a lack of signal with a lack of phylogenetic effect (Blomberg and Garland 2002),. Our statistics were not designed to test the coevolution of these traits over time, rather they look for phylogenetic covariances in the extant traits

Life history, diet, and other differences may also explain some variation in antibacterial capacity curves among mammalian species. The pace-of-life hypothesis posits that reproductive rate and longevity, among other life-history traits, should shape investment in immune defenses (Lee 2006). Although life histories are closely tied to body mass in mammals (Peters 1983; Calder 1984), species of the same mass, but with different life histories, will have different costs and benefit structures for immune defense. For example, small species that are relatively long-lived incur the higher costs of activating immune responses (Brace et al. 2017), natural antibody titers increased with incubation period in 70 species of tropical birds after accounting for body mass statistically (Lee et al. 2008), and faster-lived birds produced more of an acute-phase protein after inoculation with *E. coli* after accounting for body mass (Robinson et al. 2010). Determining if life history explains the residual variation after accounting for body mass and phylogeny is a next step towards understanding the variation in interspecific variation in antibacterial capacity, but are beyond the scope of theis current study

## Conclusion

Our findings suggest that large and small species might use different strategies to obtain the same level of protection against parasites. They have ramifications for human medicine and management of disease cycles, especially zoonoses, and a greater understanding of immune scaling could also help improve epidemiological models. A previously developed allometric framework for host-species competence for multi-host parasites found that data on interspecific allometries of immune defenses was lacking in the literature (Downs et al. 2019). Because “the rate that pathogens spread through populations is influenced by the rate of spread through individual hosts” (De Leo and Dobson 1996), we must understand better how host traits affect their propensity to maintain and transmit viable parasites (Bolzoni et al. 2008; De Leo et al. 2016).

We expect that many more important immune allometries await description. In particular, we encourage future scaling work on Order *Chiroptera*, as bats are implicated in the emergence of many zoonoses including SARS-COV2 (Han et al. 2016; Olival et al. 2017). Unfortunately, bats are rare in zoos, so few bat species were in the present analysis (but see Ruhs et al. In press). We also encourage efforts to investigate immune allometries in ectotherms, and especially vectors, as their immune defenses are more temperature-dependent than mammals (Brown et al. 2004), an important condition for a planet undergoing extensive climate change. We also hope creative approaches can be applied to scaling studies of more pathogenic microbes, ideally in more organismal contexts. Such work is challenging, as pathogenicity differs among species as well as route of infection (Taylor 1983). By continuing to merge cellular and molecular efforts with the behavioral, ecological, and evolutionary concepts, we can produce a more integrative immunology and thus more effective options to predict and manage infectious disease risk.

## Author Contributions

C.J.D., L.B.M, and K.C.K. conceptualized the study; L.A.S., S.J.O. and C.J.D., and R.B. procured samples; L.A.S., S.J.O. and C.J.D. established and validated the laboratory methodology; L.A.S., S.J.O., C.J.D., and L.B.M collected and curated the data; C.J.D. carried out the formal analysis with input from L.B.M and E.W.G; C.J.D and L.B.M drafted the manuscript with input from K.C.K; all authors contributed to writing and revision; C.J.D and L.B.M. administered and supervised the project; C.J.D., L.B.M., and R.H.Y.J acquired funding.

## Acknowledgements

We thank Birmingham Zoo, Busch Gardens Tampa Bay (Cara Martel), Cleveland Metroparks Zoo, Columbus Zoo, the Denver Zoological Foundation, Dickerson Park Zoo, Indianapolis Zoo, Lincoln Park Zoo, Louisville Zoological Garden, Maryland Zoo in Baltimore, Milwaukee County Zoo, Naples Zoo, Nashville Zoo at Grassmere, Oregon Zoo (Mitch Finegan & David Shepherdson), Sedgwick County Zoo, Smithsonian’s National Zoo and Conservation Biological Institute, Topeka Zoo & Conservation Center, Zoo Atlanta (Hayley Murphy, Stephanie Earhardt), ZooTampa at Lowry Park, and the Demas (Indiana State University), Hoekstra (Harvard University), Ophir (Cornell University), Place (Cornell University), and Monteith (University of Wyoming) labs for donating serum samples. We thank M. Amy, B. Butler, K. Carlson, E. Chinchilli, K. Clausen, H. Droke, M. Espino, A. Fernandez, M. Galan, M. Henning, N. Huizenga, K. Koller, E. Lee, O. Ogunsina, V. Pappademetriou, A. Roberts, C. Stanell, S. Travis, A. Vorrath, R. Winner, C. Yeung for help with laboratory work. N. Keith helped develop the Program R workflows required to process our data. We thank N.A. Dochtermann and J.C. Uyeda for statistical guidance, and the Martin Lab, HA Woods, J.F. Harrison, ME Sobolewski, and V Savage for discussion about an early version of the manuscript. This work was supported by State University of New York College of Environmental Science and Forestry’s Provost Office, the Levitt Center, Hamilton College’s Dean of Faculty Office, University of South Florida’s College of Public Health, and the National Science Foundation (award numbers IOS 1656551 to C.J.D. and IOS 1656618 to L.B.M).

## Data Accessibility

The datasets generated during and/or analyzed during the current study and will be deposited in Figshare after a 3-year embargo. We will consider written requests for the data prior to the end of the embargo. Lab protocols (Schoenle et al. 2020) and R code (Keith et al. 2020; Downs et al. 2021) are available on FigShare.

