## Supplementary Materials for "Large Mammals Have More Powerful Antibacterial Defenses Than Expected from Their Metabolic Rates"

**This PDF file includes:**

Supplementary Methods
Figs. S1 to S7
Tables S1 to S6
Captions for Movies S1

**Other Supplementary Materials for this manuscript include the following:**

Movies S1

### Supplementary Methods

#### *Bacterial Species*

Bacterial species used and safety approvals:

*Escherichia coli* (ATCC #8739) and *Micrococcus luteus* (ATCC # 4698) are biosafety-level 1 bacteria. *Salmonella enterica* (ATCC #13311) is a biosafety-level 2 bacterium. These strains are commonly used in comparative studies (Tieleman et al. 2005; Matson et al. 2006; Demas et al. 2011) and represent diverse bacteria types; *E. coli* and *S. enterica* are Gram-negative bacteria whereas *M. luteus* is Gram-positive to Gram-variable. For *S. enterica,* all steps of the following procedures were performed in a Biosafety cabinet (Labconco Class 2 A2 BSC). All procedures were approved by appropriate institutions at the author's home institutions.

Reconstituting purchased bacteria:

*E. coli* and *S. enterica* were purchased as frozen 1 ml aliquots in glycerol (American Type Culture Collection, Manassas, VA 20108, mini-pack *E. coli* #8734, *S. enterica* #8739). We initially thawed them in a 37 °C shaking water bath for 1-3 min. *M. luteus* was purchased as lyophilized pellets (Microbiologics 0242L). We used sterile technique to transfer a pellet to a sterile 10 ml falcon tube containing 500 µl of sterile Dulbecco's Phosphate Buffered Saline (PBS, Sigma-Aldrich #D8537). The mixture was heated in a 30-37°C water bath for 2 minutes and vortexed until the pellet was dissolved.

#### *Adapting the assay for a multispecies comparison*

Serum used for calibrations

Serum samples for calibration steps were purchased from commercial vendors, donated by zoos, or were extra samples from other studies. Samples from zoos were collected prior to our study or upon routine examination of animals. We used serum from diverse mammal species for which we were able to obtain: 1) large sample volumes (e.g., elephants, rhinos), or 2) multiple samples from the same species that we could pool to obtain enough serum for the entire plate. We obtained domestic cat (*Felis catus*), Cetacean species (dolphin or whale), cow (*Bos taurus*) and Asian elephant (*Elephas maximus*) serum samples from Ned Place at Cornell University and bighorn sheep (*Ovis canadensis*) samples were provided by Kevin Monteith at University of Wyoming. Amur and Sumatran Tigers (*Panthera tigris*), and African lion (*Panthera leo*) sera and African elephant (*Loxodonta africana*) samples were obtained from zoo partners (ZooTampa, Smithsonian's National Zoo and Conservation Biological Institute). We purchased pooled CD1 mouse, cow, and rabbit (*Oryctolagus cuniculus*) serum samples (with complement) from a commercial provider (Innovative Research Novi, MI 48377, CD1 mouse #IGMSCD1CSER, cow: #IBV-Compl, rabbit: # IGRBCSER). Use of these samples was approved by the Institutional Animal Care and Use Committees at the author's home institutions.

Step 1: Establishing growth curves

*Methods:* We aimed to harvest our bacteria for use in assaying our experimental samples while they were growing exponentially (i.e., when cultures were in the upper 50-80% of the linear part of logistic growth curves). To do so, we first needed to establish the growth curve for each type of bacterium by growing each bacterium for 15 (*E. coli* and *S. enterica*) or 33 hours (*M. luteus*). We measured growth as absorbance (optical density, OD) and compared absorbance against colony-forming units (CFU) quantified on Petri dishes. We used the same basic procedure for each bacterium and identified differences where necessary.

Purchased bacteria were reconstituted/thawed, as described previously. For each bacterium, the thawed, reconstituted bacteria mixture was then added to a 1000 ml flask containing 200 mL sterile tryptic soy broth (TSB, BD #211825 reconstituted in distilled water per manufacturer's instructions) and covered with sterilized aluminum foil. Flasks were placed in a shaking water bath at 37 °C for the duration of the growth trial. A 500 µL sample of bacteria mix was collected approximately every hour (but see details of the schedule below) and was used to quantify absorbance and CFU. To quantify absorbance, we serially diluted the bacteria mixture in TSB (raw, 1:2, 1:4, 1:8) on a 96-well plate. Each well contained 60 µl of liquid, and dilutions were run in triplicate. Each plate was shaken for 3 s, and absorbance was quantified using a Biotek Synergy HTX Multi-Mode Reader set to 300 nm. To quantify CFUs, solutions were plated on tryptic soy agar (TSA, BD #236950 reconstituted in distilled water per manufacturer's instructions) plates. We made 4-5 dilutions at each sampling period by mixing the bacteria mix with TSB. Dilutions could include 1:10, 1:100, 1:1000, 1:10,000, 1:100,000, 1:1,000,000, 1:5,000,000, and 1:10,000,000. Dilutions were chosen based on preliminary experiments (data not shown) and ensured that at least one of the dilutions would be countable (between 50 and 300 colonies per 100 mm diameter plate). Twenty-five µL of each dilution was plated in triplicate for each sampling hour. Colonies were counted following incubation in a 37°C incubator for 12-24 hours (time was dependent on the bacterium). We plotted growth curves and correlated absorbance readings and CFU in Excel.

*Results:* Data indicated that harvest of *E. coli* would be optimal at hour 6, *S. enterica* at hour 8, and *M. luteus* at hour 27 (Fig. S5). The expected number of CFUs at harvest was $10^8$-$10^9$ for all bacteria.

Step 2: Growing and cryopreserving stocks of bacteria

We used the same basic procedure to grow and cryopreserve stocks of each of the three bacteria. Purchased bacteria were reconstituted/thawed as described previously. For each bacterium separately, we added the reconstituted/thawed bacteria mixture to 200 mL of sterile TSB in a 1000 ml flask and covered it with sterilized aluminum foil. Mixtures were incubated at 37 °C on a shaker for the time determined during step 1. After incubating, we collected a 500 µL sample, then quantified absorbance using the procedure described in Step 1. We compared results to those from Step 1 to ensure that the final concentrations of bacteria were within the expected range. The bacteria mixture was portioned evenly between five 50-ml conical vials, which were centrifuged at 3000 rpm for 8 min (room temperature) to pellet bacteria. Broth was pipetted off the pellets and replaced with the equivalent volume of TSB with 10% glycerol to resuspend the pellet. One mL aliquots were transferred to sterile cryovials. Vials were cool samples at a rate of 1ºC per minute for 75 minutes using CoolCells (Corning) in a -80ºC freezer. Stock bacteria were stored at -80°C until used.

Step 3: Viability testing and quantifying CFU of stocks

One to two weeks post-cryopreservation, four to six cryovials of each bacterial species were chosen arbitrarily for viability testing. Vials were placed inside 50 ml conical vials and thawed in a 30-37°C water bath for 4-5 minutes. We serially diluted each bacterial species on a 96-well plate using the protocol described in Step 1 with additional serial dilutions of 1:16 and 1:32. We added 100 µL of sterile TSB to all wells. Covered plates were incubated at 37ºC. Absorbance at

300 nm was measured at hours 0, 8, and 24 of incubation. Before each reading, plates were shaken at 700 rpm for 1 min. We plotted growth curves to ensure that bacteria were viable and grew as expected during the 24 hr incubation.

At hour 0, we also made 1:50,000, 1:100,000 and 1:500,000 dilutions of each stock bacteria in TSB. Twenty-five µL of each dilution was plated in triplicate on TSA plates. Colonies were incubated for 24h at 37°C and then counted. We used these counts to determine the final concentration (CFU/ml) of the frozen aliquots of the stock bacteria.

*Results:* All samples of frozen stock bacteria were viable and demonstrated the expected growth after 24 hours (data not shown). The starting concentration was $10^9$ for *E. coli*, $10^9$ for S. *enterica*, $10^7$-$10^9$ for *M. luteus* (depending on the harvest).

<u>Step 4: Optimization for a multispecies comparison</u>

We aimed to optimize the antibacterial assays to compare species with potentially vastly different antibacterial ability. We used the assay developed by French and Neuman-Lee (2012) as the basis for our assay, but rather than measuring antibacterial capacity at a single bacteria/serum dilution, we quantified antibacterial capacity across a 12-point curve of serum dilutions. Briefly, for the base assay, a known amount of bacteria was mixed with serum diluted in PBS in a 96-well plate. The positive control wells contained the bacteria and volume of PBS equal to the sum of the volumes of PBS and serum in the sample wells. Positive control wells contained the volume of PBS equal to the sum of the volumes of PBS and serum sample wells. Negative control wells contained the volume of PBS equal to the sum of the volumes of bacteria, PBS, and serum sample wells. Samples and controls were plated in (at least) triplicate. Plates were covered, shaken for 1 min. at 700 rpm, and then incubated for 30 min at 37 °C. This incubation temperature is commonly used (e.g., French and Neuman-Lee 2012) and is close to the mean body temperature of mammals held in thermoneutral conditions (mean ± sd = 36.1 ± 1.9 °C, n=441, data from (White et al. 2006)). After incubation, each plate was shaken for 1 min at 700 rpm, 125 µL of TBS was added to all wells, and the plate was shaken again for 1 min at 300 rpm. Absorbance was measured at 300 nm. The first reading served as a control for the later reading taken for calculating antibacterial capacity. The covered plate was then incubated at 37 °C until growth differences became quantifiable (see below). Each plate was shaken for 1 min at 300 rpm and read again at 300 nm to obtain final measures of bacterial growth. Antibacterial capacity for each dilution of each well was calculated as $\left(1 - \frac{(sample\ end - sample\ baseline)}{(control\ start - control\ end)}\right) * 100\%$.

We optimized bacteria concentration, bacteria volume, duration of the second, longer incubation, and serum dilutions using two optimizations assays. First, we measured the growth of bacteria from stocks of different concentrations and with different starting volumes. We did not add serum so we could quantify the growth dynamics when the bacteria were grown within the parameters defined by the antibacterial assay (as opposed to in a large beaker of growth media). Second, we measured antibacterial ability of 6-9 mammalian species at 29 serum dilutions to determine the 12 dilutions that would best-capture the variation in antibacterial capacity among mammalian species.

*Methods. Optimization step 1:* In this first optimization step, we measured growth of 4, 5, 6, 8 and 10 μL of working bacteria concentrations of $10^2$, $10^3$, $10^4$, and $10^5$ CFU in the absence of serum to determine the bacteria concentration and volume and the incubation duration most appropriate for each bacterial species. Each combination was plated in triplicate on a 96-well plate. We made up working solutions of bacteria from 3 frozen aliquots of each bacterium to check for consistencies of stocks. Each stock was plated on a different 96-well plate for a total of 3 plates per bacterium. The location of each concentration-volume combination was spread randomly across plates to minimize any well-location effects. First, three frozen bacteria stocks of each bacterium were thawed as described in step 3 and 300 μL of each required working concentration was prepared in sterile PBS. The appropriate volume of the working concentrations was added to the well and PBS was added as necessary to obtain a final total volume of 10 μL. We then added 125 μL of TSB to each well as growth media. We measured absorbance at 300 nm at hour 0 following a 3s shake. Covered plates were then incubated at 37 °C. Additional readings were taken at 2 h intervals as appropriate for each bacterium (h 6-16 for *E. coli* and *S. enterica*, and h 10-22 for *M. luteus*). We graphed growth curves and visually discerned the best concentration(s) and volume(s) of bacteria in optimization step 2. When no dilution/volume combination was obviously the best in Optimization Step 1, we tested more than one in Optimization Step 2. We also visually determined the time(s) at which the population size peaked. We used this information to determine the best range of times to measure samples during optimization step 2.

*Results. Optimization step 1:* Working bacteria concentrations from $10^3$ to $10^6$ showed growth that was measurable on our plate readers at an absorbance reading of 300 nm. Starting concentrations of $10^3$ did not clearly reach maximal population size for any of the bacteria; no clear peak population size was ever reached for *M. luteus*. The best working concentration of each bacterial species were $10^4$ or $10^5$ CFU $ml^{-1}$ for *E. coli* and $10^3$ or $10^4$ CFU $ml^{-1}$ for *S. enterica* and *M. luteus*, but results were ambiguous for these concentrations so we chose to pursue these candidate concentrations in optimization step 2. Because we did not want to overwhelm the antibacterial capacity of species with relatively low antibacterial ability in our final assays, we were conservative and chose to use 2 μL of the working bacterial concentration in optimization step 2. We narrowed the long incubation ranges for the bacteria as follows: *E. coli* incubates for 10-14 hours, *S. enterica* incubates for 8-12 hours, and *M. luteus* incubates for 24-48 hours (Fig. S6).

*Methods: Optimization step 2:* The aim of this second step was to determine a series of assay conditions that would allow for multispecies comparison, especially appropriate serum dilutions, a volume and concentration of bacteria, and the length of the incubation period for bacterial growth post-antibacterial activities (i.e., the length of the second incubation). To do so, we ran 29-point dilution curves (a dilution factor of 2 from raw serum to 1:128, a dilution factor of 3 from 1:3 to 1:384, a dilution factor of 5 from 1:5 to 1:640 and a dilution factor of 3:4 to 3:64) for serum from 6-10 distinct mammalian species following the steps described previously (Step 4: Optimization for a multispecies comparison) using the serum samples describe under "Serum used for calibrations". For all three bacteria, we tested CD1 mouse, horse, cat, cow, dog, cetacean, rabbit and elephant. Additional we tested bighorn sheep for *E. coli* and *S. enterica*. For each plate, we made four serial dilutions of sera in PBS to test 29 dilutions simultaneously: a dilution factor of 2 from raw serum to 1:128, a dilution factor of 3 from 1:3 to 1:384, a dilution factor of 5 from 1:5 to 1:640 and a dilution factor of 3:4 to 3:64. Serum dilutions were plated in triplicate and the final volume of serum plus PBS was 18 ul. We also plated 6 replicates of positive and 3 replicates of negative controls. We used 2 µL of the working concentrations of each bacterium ($10^4$ or $10^5$ CFU $ml^{-1}$ for *E. coli* and $10^3$ or $10^4$ CFU $ml^{-1}$ for *S. enterica* and *M. luteus*) and incubated plates at 37 °C for various time durations, which depended on the bacterium used in the assay and were chosen from the observed growth dynamics. We measured absorbance at 300 nm at hour 0, 8, 10 and 12 for *E. coli* assays; 0, 8, 9, 10, 12 for *S. enterica* assays; and 0, 24, 26, 30, and 48 for *M. luteus* assays. We plotted the dilution curves and chose dilutions for a 12-point curve that would cover the range of antibacterial abilities exhibited by these species, which would inform the dilutions we used in the final study.

*Results. Optimization step 2:* Results of the dilution curves for 7 species against *E. coli* are depicted in Fig. S7. The final 12-point curve for each bacterium is listed in Table S5. Briefly, *E. coli* had two serial dilutions: 6 points from raw to 1:64, and 6 points from 3:4 to 3:256. *S. enterica* also had two serial dilutions: 6 points from raw to 1:64, and 6 points from 3:4 to 3:128. *M. luteus* had a single serial dilution ranging from raw serum to 1:2048. When we later used this assay for more species, we learned that our *E. coli* and *S. enterica* curves did not capture points where antibacterial capacity switched from 100% to 0%; that is, a few species had an antibacterial capacity of 100% at the lowest dilution on our 12-point dilution curve. We used serum from African lions and Amur and Sumatran tigers to build a 'super-antibacterial capacity' 12-point dilution curve. These curves included serum dilutions that were more dilute than those included on these original curve (*E. coli*: 1:4, 1:8, 1:32, 1:64, 1:128, 1:256, 3:16, 3:32, 3:64, 3:128, 3:256, 3:512; *S. enterica*: 1:16, 1:32, 1:64, 1:128, 1:256, 1:512, 3:4, 3:8, 3:16, 3:32, 3:64, 3:128). The cow serum dilutions that captured the shift from 100% antibacterial capacity to 0% differed by bacterial species and were 1:32, 1:64, 1:128, 1:256 for *E. coli*; 1:16, 1:32, 1:64, 1:128 for *S. enterica*; and 1;20, 1:80, 1:320, 1:1280 for *M. luteus.* The best bacterial challenge for the samples was 2 µl of $10^4$ CFU $ml^{-1}$ for all bacterial species. The best long incubation duration was 12 h for *E. coli*, 10 h for *S. enterica*, and 48 h for *M. luteus* at an incubation temperature of 37 °C. The key parameters of the optimized antibacterial assay against *E. coli*, *S. enterica*, and *M. luteus* are summarized in Table S5. Protocols for antibacterial capacity dilution corves against all three bacterial species are available in FigShare (Schoenle et al. 2020).

**Data analysis: Phylogenetic univariate models.**

*Methods.* To check for effects of mammalian phylogeny on antibacterial capacity, we constructed a separate phylogenetic univariate mixed model for each curve parameter for each bacterium. We included body mass and all other curve parameters as fixed effects then fit models using `MCMCglmm` (Hadfield 2010; Hadfield and Nakagawa 2010). The phylogenetic covariance matrix for this analysis was estimated using a phylogenetic tree constructed with NCBI molecular data and phyloT (Fig. S4) (Letunic 2015). All mixed models were fit using a weak inverse-Gamma prior with shape and scale parameters set to 0.01 for the random effect of phylogenetic variance. Default priors for all other fixed effects were used. Model chains were run for $7.8 \times 10^5$ iterations, with an 180,000-iteration burn-in and a 600-iteration thinning interval. We estimated Pagel's lambda as a measure of how much of the total observed variation was explained by phylogeny (Housworth et al. 2004).

*Results.* Phylogenetic univariate models supported scaling coefficients of zero and that phylogeny explained <19% of the variation in antibacterial curve parameters (see main text for more details). Individual body mass was not a significant predictor of any of the curve parameters for antibacterial capacity against any of the bacterial species. At least one of the curve parameters was a significant fixed effect each curve parameter (response variable) for antibacterial capacity against *E. coli* (Table S6), *S. enterica* (Table S7), and *M. luteus* (Table S8).


Demas, G. E., D. A. Zysling, B. R. Beechler, M. P. Muehlenbein, and S. S. French. 2011. Beyond phytohaemagglutinin: assessing vertebrate immune function across ecological contexts: Assessing vertebrate immune function across ecological contexts. Journal of Animal Ecology 80:710–730.

French, S. S., and L. A. Neuman-Lee. 2012. Improved ex vivo method for microbiocidal activity across vertebrate species. Biology Open 1:482–487.

Hadfield, J. D. 2010. MCMC Methods for Multi-Response Generalized Linear Mixed Models: The **MCMCglmm** *R* Package. Journal of Statistical Software 33.

Hadfield, J. D., and S. Nakagawa. 2010. General quantitative genetic methods for comparative biology: phylogenies, taxonomies and multi-trait models for continuous and categorical characters. Journal of Evolutionary Biology 23:494–508.

Housworth, E. A., E. P. Martins, and M. Lynch. 2004. The Phylogenetic Mixed Model. The American Naturalist 163:84–96.

Letunic, I. 2015. phyloT: phylogenetic tree generator.

Matson, K. D., A. A. Cohen, K. C. Klasing, R. E. Ricklefs, and A. Scheuerlein. 2006. No simple answers for ecological immunology: relationships among immune indices at the individual level break down at the species level in waterfowl. Proceedings of the Royal Society B: Biological Sciences 273:815–822.

Schoenle, L. A., L. B. Martin, and C. J. Downs. 2020. Protocols for 12-dilution antibacterial capacity curves for interspecific comparisons. Figshare. 10.6084/m9.figshare.12501149

Tieleman, B. I., J. B. Williams, R. E. Ricklefs, and K. C. Klasing. 2005. Constitutive innate immunity is a component of the pace-of-life syndrome in tropical birds. Proceedings of the Royal Society B: Biological Sciences 272:1715–1720.

White, C. R., N. F. Phillips, and R. S. Seymour. 2006. The scaling and temperature dependence of vertebrate metabolism. Biology Letters 2:125–127.

**Fig. S1.**

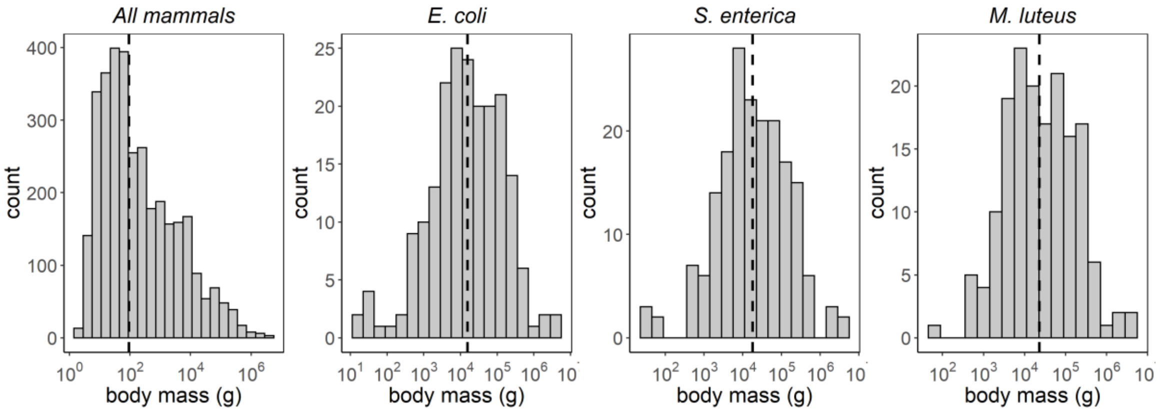


Histogram of the distribution of body mass for all extant mammals (# species = 3350, median body mass = 92.5 g; data from Smith et al. 2003; A) and those used to measure antibacterial capacity against *Escherichia coli* (# species = 199, median body mass = 15,610 g; B), *Salmonella enterica* (# species =186, median body mass = 18,062 g; C), and *Micrococcus luteus* (# species = 164, median body mass = 22,847 g; D). Dashed lines depict median body size.

**Fig. S2.**

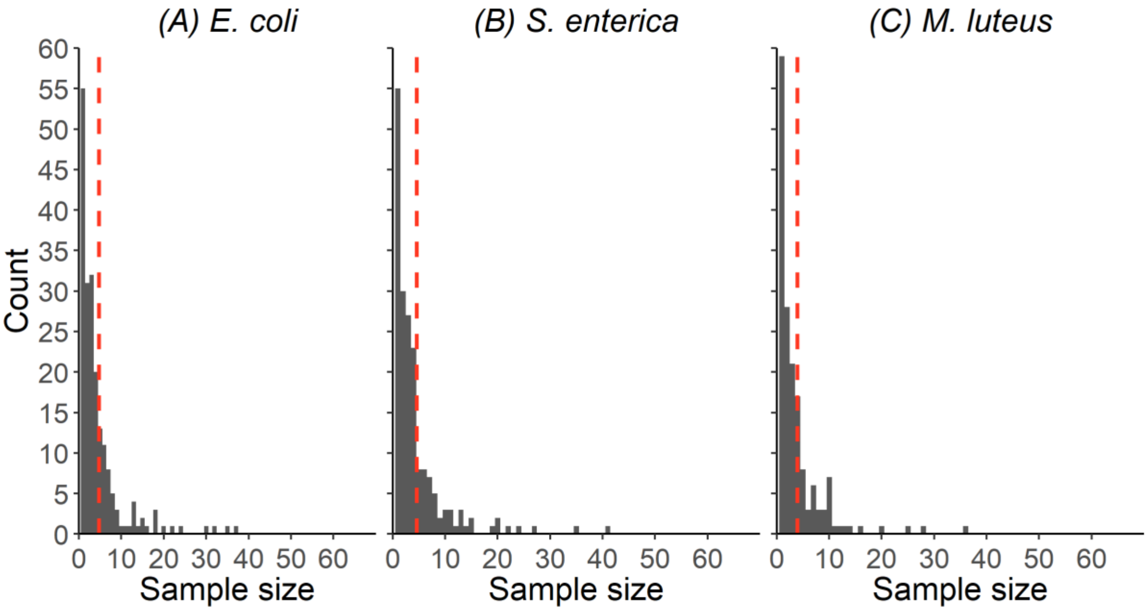


Histogram of the number of samples for each species analyzed for antibacterial capacity against *Escherichia coli* (A), *Salmonella enterica* (B), and *Micrococcus luteus* (C). The red dashed lines denote mean sample size (n=4.7 for A, 4.6 for B, and 3.9 for C).

**Fig. S3.**

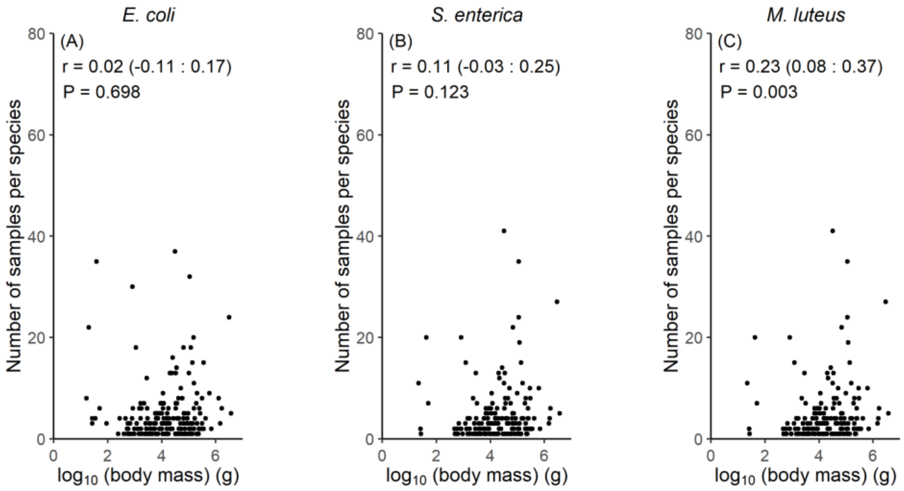


More samples were used for larger species for one of three bacteria species *Micrococcus luteus* (C), but not *Escherichia. coli* (A) or S*almonella enterica* (B). Although, this correlations was weak.

**Fig. S4.**

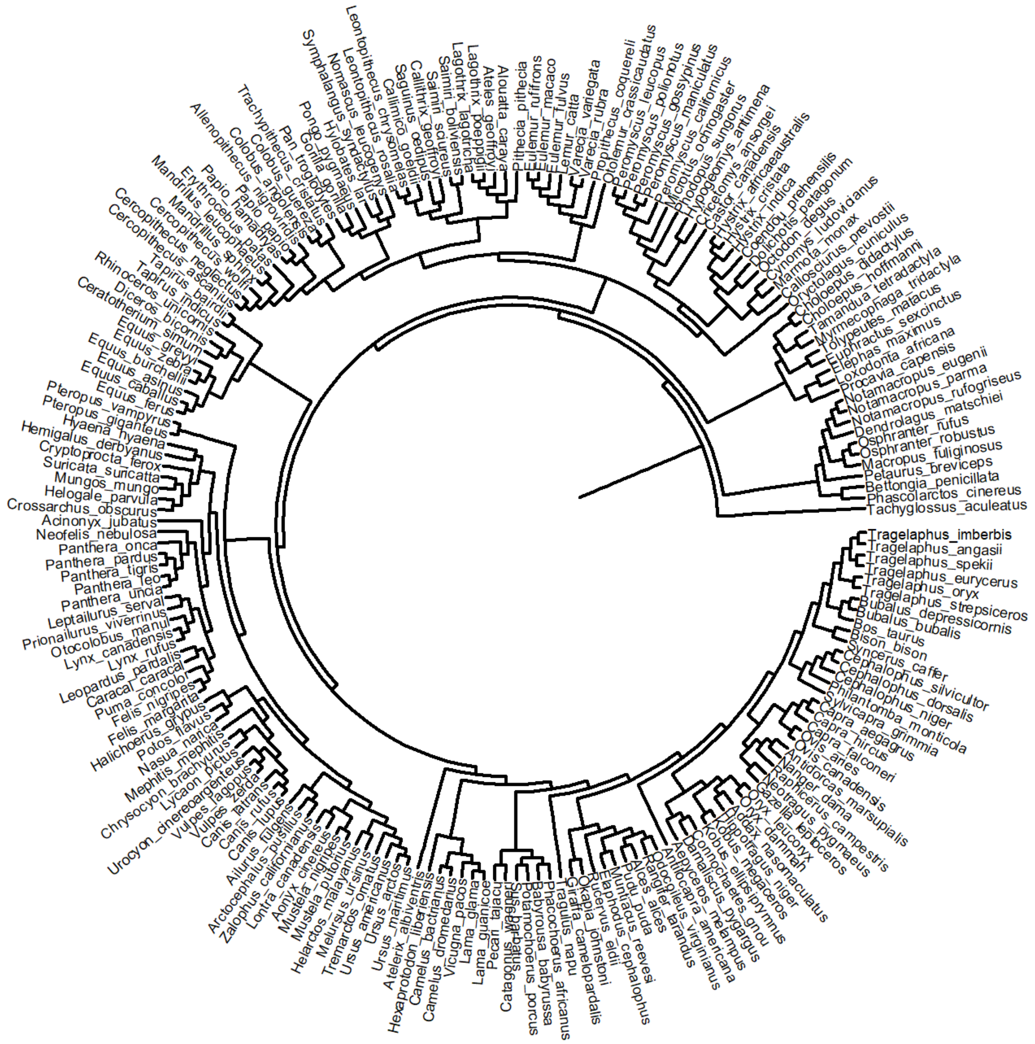


Phylogenetic tree (not time rooted) depicting the relationship among the 199 species uses in this study. Nodes represent hypothesized shared ancestors.

**Fig. S5.**

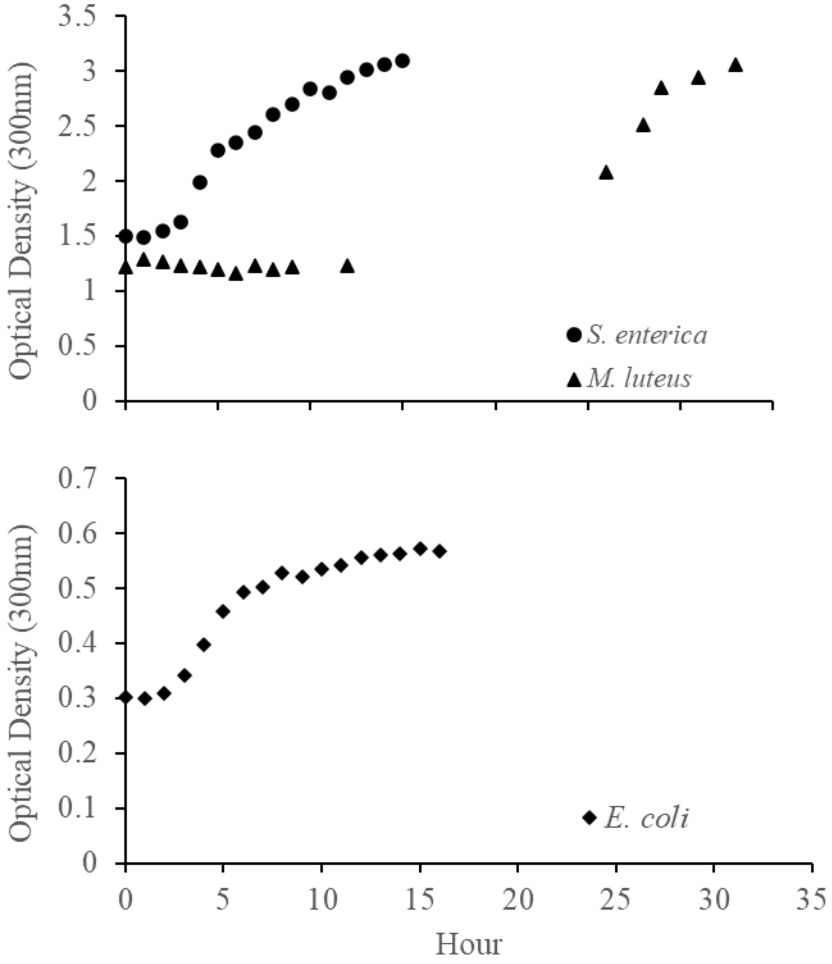


Growth curves of *Salmonella enterica* (A), *Micrococcus luteus* (A), and *Escherichia coli* (B). Each point represents the mean calculated from 3 replicate samples. We were unable to collect data for all time points because of staffing.

**Fig. S6.**

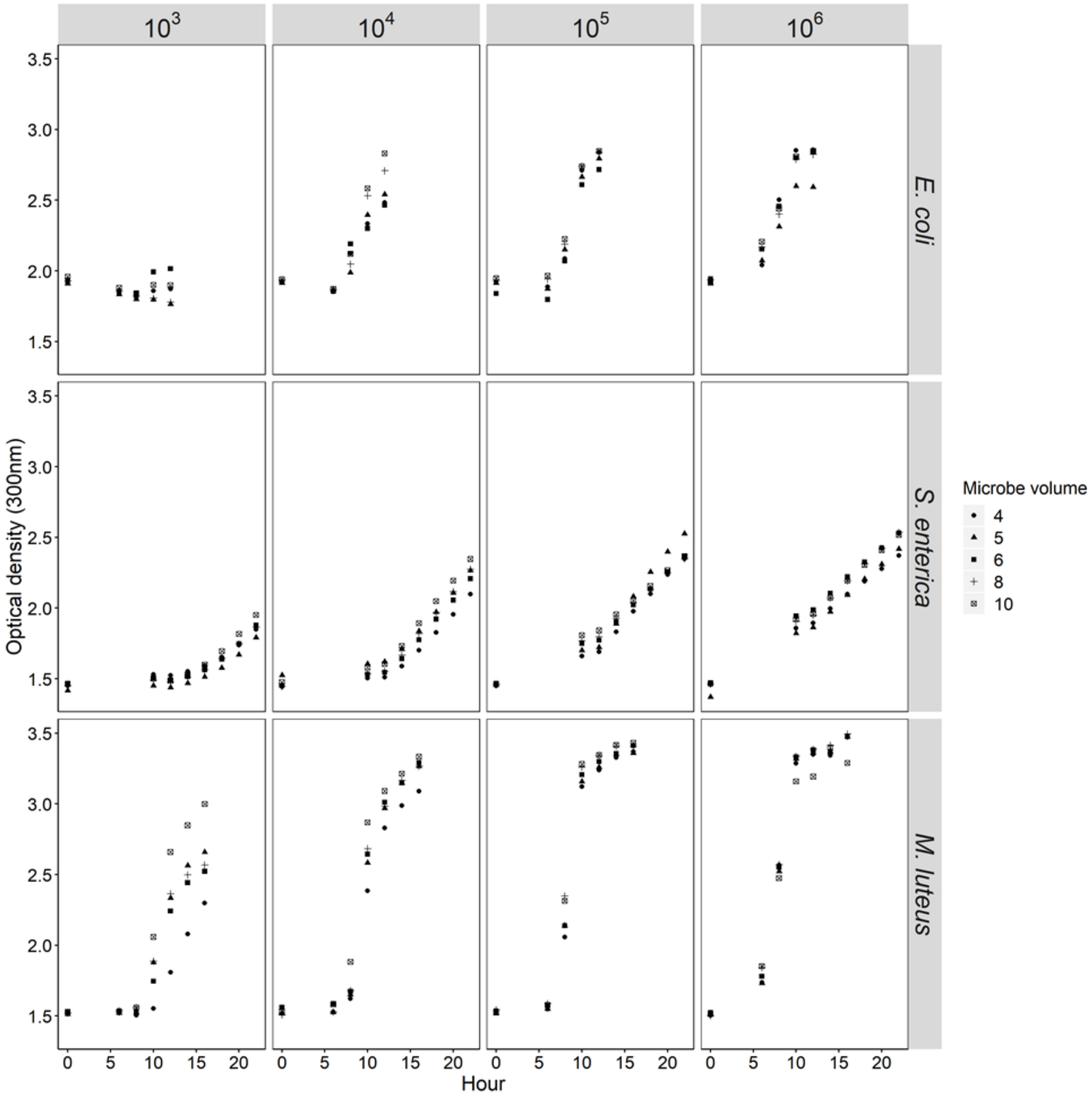


Growth curves for *Escherichia. coli*, *Salmonella enterica*, and *Micrococcus luteus* from Optimization step 1.

**Fig. S7.**

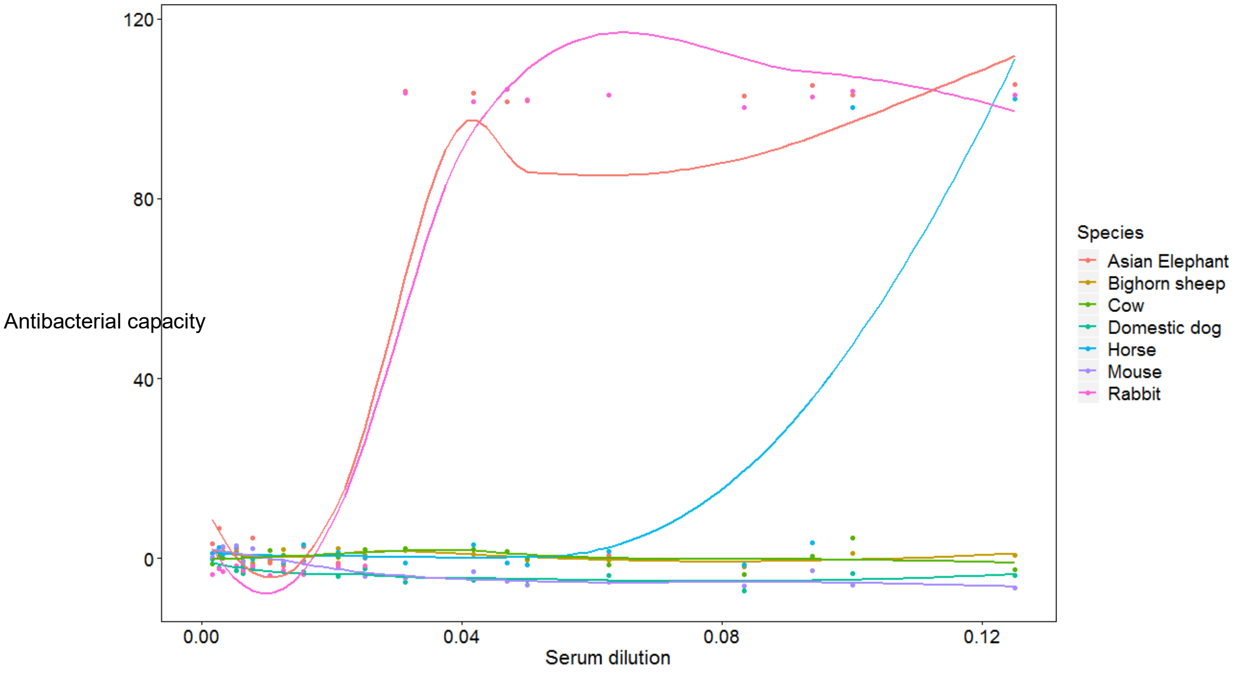


Antibacterial capacity against *E. coli* of 27 dilutions of serum for 7 mammalian species. Lines are fit using spline. This graph is an example of the types of curves we antibacterial capacity curves obtained during the calibrations of the assay. These samples were used to in the final analyses because they did not meet our inclusion criteria.

**Table S1.** Complete interspecific (co)variances of antibacterial capacity curve parameters and body mass. $\text{Inflection}_{\text{antibacterial capacity}}$, top asymptote, and bottom asymptote were+1, $\log_{10}$-transformed. Body mass, slope and $\text{inflection}_{\text{dilution}}$ were $\log_{10}$-transformed.

| | Top asymptote | Slope | $\text{Inflection}_{\text{dilution}}$ | Asymptote coefficient | $\text{Inflection}_{\text{antibacterial capacity}}$ | Bottom asymptote | Body mass |
|---|---|---|---|---|---|---|---|
| ***E. coli*** | | | | | | | |
| Top asymptote | 0.003<br>($9.7 \times 10^{-4}$ : 0.010) | -0.649<br>(-2.408 : 0.676) | 0.033<br>(0.013 : 0.070) | 0.003<br>(-0.003 : 0.014) | 0.003<br>($4.8 \times 10^{-4}$ : 0.008) | -0.002<br>(-0.005 : $6.0 \times 10^{-4}$) | -0.002<br>(-0.036 : 0.035) |
| Slope | | 847.4<br>(445.9 : 1983.5) | -12.2<br>(-27.6 : -2.3) | 1.40<br>(-2.33 : 5.22) | -1.005<br>(-2.666 : 0.019) | -1.04<br>(-2.7 : -0.19) | 9.50<br>(-4.3 : 26.3) |
| $\text{Inflection}_{\text{dilution}}$ | | | 0.431<br>(0.233 : 2.00) | 0.020<br>(-0.093 : 0.097) | 0.030<br>(0.008 : 0.060) | -0.012<br>(-0.069 : 0.008) | -0.262<br>(-0.723 : 0.360) |
| Asymptote coefficient | | | | 0.013<br>(0.005 : 0.139) | 0.001<br>(-0.006 : 0.010) | -0.005<br>(-0.014 : 0.008) | -0.022<br>(-0.345 : 0.178) |
| $\text{Inflection}_{\text{antibacterial capacity}}$ | | | | | 0.002<br>($6.7 \times 10^{-4}$ : 0.106) | $-5.3 \times 10^{-4}$<br>(-0.003 : 0.005) | -0.006<br>(-0.284 : 0.036) |
| Bottom asymptote | | | | | | 0.004<br>(0.003 : $2.8 \times 10^{208}$) | -0.002<br>($-1.3 \times 10^{78}$ : $2.1 \times 10^{95}$) |
| Body Mass | | | | | | | 2.49<br>(3.60 : $7.5 \times 10^{20}$) |
| ***S. enterica*** | | | | | | | |
| Slope | N/A* | 188.8<br>(45.8 : 774.5) | -8.65<br>(-22.32 : -1.20) | 0.293<br>(-1.648 : 2.173) | -0.026<br>(-1.703 : 1.852) | 0.318<br>(-0.082 : 1.088) | -0.394<br>(-11.9 : 12.6) |
| $\text{Inflection}_{\text{dilution}}$ | | | 0.594<br>(0.300 : 2.373) | 0.009<br>(-0.087 : 0.129) | 0.041<br>(-0.003 : 0.193) | -0.011<br>(-0.042 : 0.015) | -0.126<br>(-0.747 : 0.465) |
| Asymptote coefficient | | | | 0.010<br>(0.004 : 0.097) | 0.005<br>(-0.009 : 0.041) | 0.002<br>(-0.002 : 0.014) | -0.107<br>(-0.341 : 0.033) |
| $\text{Inflection}_{\text{antibacterial capacity}}$ | | | | | 0.012<br>(0.006 : 0.148) | 0.002<br>($-2.6 \times 10^{-4}$ : 0.023) | 0.009<br>(-0.146 : 0.445) |
| Bottom asymptote | | | | | | 0.001<br>($7.9 \times 10^{-4}$ : 0.090) | -0.009<br>(-0.409 : 0.046) |
| Body Mass | | | | | | | 2.31<br>(2.61 : $2.9 \times 10^{19}$) |
| ***M. luteus*** | | | | | | | |
| Top asymptote | 0.015<br>(0.004 : 0.060) | 0.332<br>(-1.78 : 2.39) | 0.096<br>(-0.004 : 0.357) | 0.009<br>(-0.012 : 0.035) | 0.011<br>(0.001 : 0.040) | 0.007<br>(-0.003 : 0.022) | -0.024<br>(-0.113 : 0.064) |
| Slope | | 174.0<br>(88.8 : 479.3) | -3.563<br>(-20.06 : 10.38) | -0.087<br>(-2.39 : 1.52) | 0.360<br>(-1.09 : 1.98) | 0.133<br>(-0.822 : 1.134) | -11.0<br>(-22.8 : -2.6) |
| $\text{Inflection}_{\text{dilution}}$ | | | 1.35<br>(0.496 : 4.158) | 0.104<br>(-0.010 : 0.322) | 0.072<br>(-0.006 : 0.263) | 0.060<br>(-0.031 : 0.184) | -0.103<br>(-0.954 : 0.947) |
| Asymptote coefficient | | | | 0.021<br>(0.008 : 0.200) | 0.005<br>(-0.018 : 0.020) | 0.009<br>(-0.001 : 0.047) | -0.022<br>(-0.390 : 0.156) |
| $\text{Inflection}_{\text{antibacterial capacity}}$ | | | | | 0.009<br>(0.001 : 14.7) | 0.005<br>(-0.090 : 0.015) | -0.023<br>(-0.087 : 4.47) |

| | | |
|---|---|---|
| Bottom asymptote | 0.005 (0.003 : 6.2 $\times 10^{64}$) | -0.041 (-2.0 $\times 10^{33}$ : 3.61) |
| Body Mass | | 2.31 (3.06 : 8.9 $\times 10^{43}$) |

* Top asymptote was not included in the analysis of antibacterial capacity against *S. enterica* because it was strong correlated with $\text{Inflection}_{\text{antibacterial capacity}}$ ($r$ = 0.998)

**Table S2**. Complete intraspecific (co)variances of antibacterial capacity curve parameters and body mass. $Inflection_{antibacterial\ capacity}$, top asymptote, and bottom asymptote were+1, $log_{10}$-transformed. Body mass, slope and $inflection_{dilution}$ were $log_{10}$-transformed.

| ***E. coli*** | Top asymptote | Slope | $Inflection_{dilution}$ | Asymptote coefficient | $Inflection_{antibacterial\ capacity}$ | Bottom asymptote | Body mass |
|---|---|---|---|---|---|---|---|
| Top asymptote | 0.023 (0.021 : 0.0256) | -0.841 (-1.322 : -0.355) | 0.044 (0.031 : 0.058) | -0.011 (-0.017 : -0.006) | 0.019 (0.019 : 0.022) | $1.3 \times 10^{-4}$ ($-8.8 \times 10^{-5}$ : 0.001) | 0.002 ($-7.2 \times 10^{-4}$ : 0.004) |
| Slope | | 2172.4 (1974.6 : 2386.9) | -8.94 (-13.10 : -4.96) | -0.295 (-2.085 : 1.469) | -0.862 (-1.317 : -0.420) | -0.593 (-0.902 : -0.287) | -0.526 (-1.287 : 0.205) |
| $Inflection_{dilution}$ | | | 1.704 (1.55 : 1.87) | -0.080 (-0.129 : -0.031) | 0.042 (0.029 : 0.055) | 0.004 (-0.005 : 0.012) | -0.007 (-0.028 : 0.013) |
| Asymptote coefficient | | | | 0.322 (0.294 : 0.353) | -0.018 (-0.024 : -0.013) | 0.001 (-0.002 : 0.005) | -0.001 (-0.010 : 0.008) |
| $Inflection_{antibacterial\ capacity}$ | | | | | 0.021 (0.019 : 0.023) | 0.006 (0.005 : 0.007) | 0.001 (-0.001 : 0.004) |
| Bottom asymptote | | | | | | 0.010 (0.009 : 0.011) | $-1.4 \times 10^{-4}$ (-0.002 : 0.002) |
| Body Mass | | | | | | | 0.052 (0.047 : 0.058) |
| ***S. enterica*** | | | | | | | |
| Slope | N/A* | 3265.7 (2959.2 : 3607.2) | 1.327 (-1.504 : 4.145) | -6.05 (-8.06 : -4.04) | 0.438 (-0.218 : 1.085) | 0.086 (-0.211 : 0.390) | -0.062 (-1.071 : 0.923) |
| $Inflection_{dilution}$ | | | 0.500 (0.451 : 0.556) | -0.091 (-0.119 : -0.065) | 0.051 (0.042 : 0.061) | -0.003 (-0.007 : $8.2 \times 10^{-4}$) | -0.004 (-0.017 : 0.009) |
| Asymptote coefficient | | | | 0.255 (0.232 : 0.281) | -0.018 (-0.024 : -0.012) | 0.006 (0.003:0.009) | -0.010 (-0.020 : $-9.3\times 10^{-4}$) |
| $Inflection_{antibacterial\ capacity}$ | | | | | 0.028 (0.025 : 0.031) | $-2.3 \times 10^{-4}$ (-0.001 : $7.8\times 10^{-4}$) | -0.002 (-0.005 : 0.001) |
| Bottom asymptote | | | | | | 0.005 (0.005 : 0.006) | $-1.2 \times 10^{-4}$ (-0.002 : 0.001) |
| Body Mass | | | | | | | 0.060 (0.054 : 0.067) |
| ***M. luteus*** | | | | | | | |
| Top asymptote | 0.045 (0.040 : 0.051) | -0.479 (-1.001 : 0.031) | 0.277 (0.227 : 0.331) | -0.005 (-0.014 : 0.004) | 0.034 (0.029 : 0.038) | -0.001 (-0.005 : 0.003) | 0.002 (-0.003 : 0.007) |
| Slope | | 849.1 (760.2 : 952.8) | 2.00 (-4.08 : 8.32) | -3.65 (-4.92 : -2.43) | 0.158 (-0.252 : 0.577) | 0.211 (-0.288 : 0.691) | 0.127 (-0.493 : 0.748) |
| $Inflection_{dilution}$ | | | 6.87 (6.13 : 7.74) | -0.255 (-0.370:-0.144) | 0.233 (0.192 : 0.277) | -0.003 (-0.050 : 0.044) | -0.010 (-0.066 : 0.049) |
| Asymptote coefficient | | | | 0.281 (0.252 : 0.315) | -0.017 (-0.024 : -0.009) | 0.0152 (0.007 : 0.024) | $1.9 \times 10^{-4}$ (-0.011 : 0.012) |
| $Inflection_{antibacterial\ capacity}$ | | | | | 0.030 (0.025 : 0.033) | 0.001 (-0.002 : 0.005) | 0.001 (-0.002 : 0.006) |
| Bottom asymptote | | | | | | 0.041 (0.037 : 0.047) | -0.003 (-0.007 : 0.002) |

| | |
|---|---|
| Body Mass | 0.060<br>(0.053 : 0.068) |

* Top asymptote was not included in the analysis of antibacterial capacity against *S. enterica* because it was strong correlated with $\text{Inflection}_{\text{antibacterial capacity}}$ ($r = 0.998$)

**Table S3.** Estimated scaling coefficients (mean ± 95% CI) for curve parameters of antibacterial capacity against *E. coli, S. enterica, M. luteus* from phylogenetic univariate mixed models. These models included a correlation matrix derived from a phylogenetic tree as a random effect to account for phylogeny. Estimates in bold have 95% CI intervals that do not overlap 0.

| | Estimated of scaling coefficient by dataset | | |
|---|---|---|---|
| **Antibacterial capacity curve parameter** | ***E. coli*** | ***S. enterica*** | ***M. luteus*** |
| Top asymptote | 0.003 (-0.003, 0.009) | $3.5 \times 10^{-4}$ (-0.007, 0.009) | -0.003 (-0.015, 0.009) |
| Slope | 0.62 (-3.18, 4.38) | 0.29 (-15.5, 15.9) | **-4.30 (-8.34, -0.30)** |
| $\text{Inflection}_{\text{dilution}}$ | -0.053 (-0.139, 0.0417) | $7.4 \times 10^{-4}$ (-0.097, 0.103) | -0.011 (-0.270, 0.262) |
| $\text{Inflection}_{\text{antibacterial capacity}}$ | -0.002 (-0.007, 0.003) | $-8.6 \times 10^{-5}$ (-0.008, 0.007) | 0.002 (-0.007, 0.013) |
| Bottom asymptote | 0.001 (-0.007, 0.008) | -0.003 (-0.016, 0.008) | -0.029 (-0.062, 0.008) |

**Table S4.** Lambda estimate from phylogenetic univariate models.

| | ***E. coli*** | ***S. enterica*** | ***M. luteus*** |
|---|---|---|---|
| Inflection$_{dilution}$ | 0.016* | 0.116 | 0.010 |
| | (0.002 : 0.035) | (0.056, 0.208) | ($3.2 \times 10^{-04}$ 0.033) |
| Inflection$_{antibacterial\ capacity}$ | 0.167 | 0.171 | 0.182 |
| | (0.111 : 0.226) | (0.111, 0.232) | (0.111 : 0.255) |
| | 0.158 | 0.155 | 0.157 |
| Top asymptote | (0.101 : 0.210) | (0.105, 0.210) | (0.093 : 0.216) |
| Bottom asymptote | 0.145 | 0.136 | 0.156 |
| | (0.072 : 0.218) | (0.069, 0.207) | (0.060 : 0.261) |
| Slope | 0.037 | 0.005 | 0.117 |
| | ($1.4 \times 10^{-06}$ : 0.080) | $8.9\times 10^{-07}$, 0.030) | (0.032 : 0.208) |

*mean and 95% CI

**Table S5.**

Key parameters of the optimized antibacterial assay conditions against *Escherichia coli*, *Salmonella enterica*, and *Micrococcus luteus.*

| Bacterial species | Dilution of working bacteria | Volume of bacteria added to wells | Serum dilutions | Final volume of serum dilutions | Cow serum dilutions | Duration of long incubation | Incubation temperature (°C) |
|---|---|---|---|---|---|---|---|
| *Escherichia coli* | $10^4$ | 2ul | Raw, 1:2, 1:4, 1:8, 1:32, 1:64, 3:4, 3:8, 3:32, 3:64, 3:128, 3:256 | 18ul | 1:32, 1:64, 1:128, 1:256 | 12 | 37 |
| *Salmonella enterica* | $10^4$ | 2ul | Raw, 1:2, 1:4, 1:8, 1:32, 1:64, 3:4, 3:8, 3:16, 3:32, 3:64, 3:128 | 18ul | 1:16, 1:32, 1:64, 1:128 | 10 | 37 |
| *Micrococcus luteus* | $10^4$ | 2ul | Raw, 1:2, 1:4, 1:8, 1:16, 1:32, 1:64, 1:128, 1:256, 1:512, 1:1024, 1:2048 | 18ul | 1;20, 1:80, 1:320, 1:1280 | 48 | 37 |
| *Escherichia coli super-antibacterial capacity* | $10^4$ | 2ul | 1:4, 1:8, 1:32, 1:64, 1:128, 1:256, 3:16, 3:32, 3:64, 3:128, 3:256, 3:512 | 18ul | 1:32, 1:64, 1:128, 1:256 | 12 | 37 |
| *Salmonella enterica super-antibacterial capacity* | $10^4$ | 2 ul | 1:16, 1:32, 1:64, 1:128, 1:256, 1:512, 3:4, 3:8, 3:16, 3:32, 3:64, 3:128 | 18ul | 1:16, 1:32, 1:64, 1:128 | 10 | 37 |

**Table S6.** Posterior means, 95% credible interval (CI), effective sample, and P-values for fixed effect from phylogenetic univariate mixed models. Each model included a single parameter from the antibacterial capacity curve against *E. coli*. $Inflection_{antibacterial\ capacity}$, top asymptote, and bottom asymptote were +1, $log_{10}$-transformed and body mass and slope were $log_{10}$-transformed prior to analyses.

| Response Variable | Fixed effect | Posterior mean | 95% CI Lower | 95% CI Upper | Effective sample | pMCMC |
|---|---|---|---|---|---|---|
| $Inflection_{dilution}$ | **Intercept** | **-0.796** | **-1.294** | **-0.251** | **1000** | **0.002** |
| | Species mean body mass | -0.053 | -0.140 | 0.042 | 1000 | 0.254 |
| | Individual body mass[a] | -0.209 | -0.584 | 0.174 | 1000 | 0.308 |
| | **$Inflection_{antibacterial\ capacity}$** | **7.60** | **4.59** | **10.7** | **1000** | **<0.001** |
| | **Bottom asymptote** | **-2.28** | **-3.90** | **-0.630** | **1000** | **0.008** |
| | **Top asymptote** | **-4.47** | **-7.23** | **-1.98** | **1000** | **0.004** |
| | **Slope** | **-0.004** | **-0.005** | **-0.002** | **1633** | **<0.001** |
| | Asymptote coefficient | 0.001 | $-7.3 \times 10^{-4}$ | 0.003 | 1560 | 0.216 |
| Top asymptote | **Intercept** | **0.099** | **0.040** | **1.24** | **997.2** | **<0.001** |
| | Species mean body mass | 0.003 | -0.003 | 0.009 | 1000 | 0.322 |
| | Individual body mass | 0.003 | -0.006 | 0.013 | 1000 | 0.51 |
| | **$Inflection_{dilution}$** | **-0.002** | **-0.004** | **$-4.6 \times 10^{-4}$** | **960.3** | **0.02** |
| | **$Inflection_{antibacterial\ capacity}$** | **1.16** | **1.14** | **1.17** | **854.4** | **<0.001** |
| | **Bottom asymptote** | **-4.80** | **-5.14** | **-4.50** | **1000.3** | **<0.001** |
| | **Slope** | **$-1.2 \times 10^{-4}$** | **$-1.6 \times 10^{-4}$** | **$-8.1 \times 10^{-5}$** | **1000** | **<0.001** |
| | **Asymptote coefficient** | **$2.6 \times 10^{-4}$** | **$2.2 \times 10^{-4}$** | **$3.1 \times 10^{-4}$** | **1000** | **<0.001** |
| Slope | **Intercept** | **67.8** | **48.1** | **87.1** | **1000** | **<0.001** |
| | Species mean body mass | 0.619 | -3.18 | 4.38 | 1000 | 0.712 |
| | Individual body mass | -7.76 | -24.3 | 8.36 | 1000 | 0.354 |
| | **Top asymptote** | **-273.7** | **-381.1** | **-177.7** | **1000** | **<0.001** |
| | **$Inflection_{dilution}$** | **-6.39** | **-9.32** | **-3.73** | **1000** | **<0.001** |
| | **$Inflection_{antibacterial\ capacity}$** | **295.7** | **181.5** | **428.0** | **1000** | **<0.001** |
| | **Bottom asymptote** | **-284.2** | **-351.3** | **-218.4** | **1000** | **<0.001** |

| | | | | | | |
|---|---|---|---|---|---|---|
| | **Asymptote coefficient** | **0.184** | **0.099** | **0.266** | **1000** | **<0.001** |
| Bottom asymptote | **Intercept** | **0.106** | **0.045** | **0.164** | **1000** | **<0.001** |
| | Species mean body mass | 0.001 | -0.007 | 0.008 | 1155 | 0.802 |
| | Individual body mass | -0.001 | -0.014 | 0.012 | 1000 | 0.914 |
| | $\text{Inflection}_{\text{dilution}}$ | -0.002 | -0.004 | 0.001 | 1000 | 0.22 |
| | **$\text{Inflection}_{\text{antibacterial capacity}}$** | **1.244** | **1.16** | **1.32** | **1000** | **<0.001** |
| | **Slope** | **$-2.3 \times 10^{-4}$** | **$-2.9 \times 10^{-4}$** | **$-1.7 \times 10^{-4}$** | **1048** | **<0.001** |
| | **Top asymptote** | **-1.04** | **-1.10** | **-0.964** | **1000** | **<0.001** |
| | **Asymptote coefficient** | **$3.1 \times 10^{-4}$** | **$2.4 \times 10^{-4}$** | **$3.9\times 10^{-4}$** | **1000** | **<0.001** |
| $\text{Inflection}_{\text{antibacterial capacity}}$ | **Intercept** | **-0.063** | **-0.910** | **-0.024** | **1093.8** | **0.002** |
| | Species mean body mass | -0.002 | -0.007 | 0.003 | 1000 | 0.382 |
| | Individual body mass | -0.0012 | -0.010 | 0.006 | 836.1 | 0.682 |
| | **$\text{Inflection}_{\text{dilution}}$** | **0.002** | **$9.3 \times 10^{-4}$** | **0.004** | **1000** | **<0.001** |
| | **Slope** | **$9.4 \times 10^{-5}$** | **$5.9 \times 10^{-5}$** | **$1.3 \times 10^{-4}$** | **1000** | **<0.001** |
| | **Bottom asymptote** | **4.12** | **3.83** | **4.37** | **917.2** | **<0.001** |
| | **Top asymptote** | **8.27** | **8.15** | **8.38** | **1000** | **<0.001** |
| | **Asymptote coefficient** | **$-2.4 \times 10^{-4}$** | **$-2.8 \times 10^{-4}$** | **$-1.9 \times 10^{-4}$** | **1117.5** | **<0.001** |

[a] individual deviation from the species mean

**Table S7.** Posterior means, 95% credible interval (CI), effective sample, and P-values for fixed effect from phylogenetic univariate mixed models. Each model included a single parameter from the antibacterial capacity curve against *S. enterica*. $\text{Inflection}_{\text{antibacterial capacity}}$, top asymptote, and bottom asymptote were +1, $\log_{10}$-transformed and body mass and slope were $\log_{10}$-transformed prior to analyses.

| Response Variable | Fixed effect | Posterior mean | 95% CI Lower | 95% CI Upper | Effective sample | pMCMC |
|---|---|---|---|---|---|---|
| $\text{Inflection}_{\text{dilution}}$ | **Intercept** | **-1.30** | **-1.89** | **-0.700** | **1000** | **<0.001** |
| | Species mean body mass | 0.001 | -0.097 | 0.104 | 1000 | 0.976 |
| | Individual body mass[a] | -0.024 | -0.197 | 0.163 | 1000 | 0.81 |
| | **$\text{Inflection}_{\text{antibacterial capacity}}$** | **3.04** | **2.05** | **4.22** | **1228** | **<0.001** |
| | Bottom asymptote | -0.017 | -0.591 | 0.566 | 1000 | 0.956 |
| | **Top asymptote** | **-0.982** | **-1.91** | **-0.076** | **1000** | **0.038** |
| | **Slope** | **-0.001** | **-0.002** | **$2.8 \times 10^{-4}$** | **1000** | **0.006** |
| | Asymptote coefficient | 0.000 | -0.001 | 0.002 | 1000 | 0.42 |
| Top asymptote | **Intercept** | **0.074** | **0.026** | **0.126** | **1000** | **0.002** |
| | Species mean body mass | 0.000 | -0.007 | 0.009 | 1000 | 0.932 |
| | Individual body mass | 0.004 | -0.012 | 0.018 | 1000 | 0.614 |
| | **$\text{Inflection}_{\text{dilution}}$** | **-0.005** | **-0.011** | **-0.001** | **1000** | **0.04** |
| | **$\text{Inflection}_{\text{antibacterial capacity}}$** | **1.17** | **1.14** | **1.19** | **1096** | **<0.001** |
| | **Bottom asymptote** | **0.103** | **0.056** | **0.146** | **1000** | **<0.001** |
| | **Slope** | **$-4.2\times 10^{-4}$** | **$-4.7 \times 10^{-4}$** | **$-3.6 \times 10^{-4}$** | **1000** | **<0.001** |
| | **Asymptote coefficient** | **$2.5 \times 10^{-4}$** | **$01.6\times 10^{-4}$** | **$3.4 \times 10^{-4}$** | **1000** | **<0.001** |
| Slope | **Intercept** | **56.3** | **34.7** | **75.1** | **1000** | **<0.001** |
| | Species mean body mass | -0.920 | -4.99 | 2.69 | 1000 | 0.662 |
| | Individual body mass | 0.292 | -15.5 | 15.9 | 1000 | 0.99 |
| | **$\text{Inflection}_{\text{dilution}}$** | **-10.3** | **-15.3** | **-5.12** | **1000** | **<0.001** |
| | **$\text{Inflection}_{\text{antibacterial capacity}}$** | **604.3** | **516.8** | **685.50** | **1000** | **<0.001** |
| | **Bottom asymptote** | **63.6** | **11.9** | **114.1** | **908.4** | **0.024** |
| | **Top asymptote** | **-503.3** | **-574.2** | **-438.4** | **1000** | **<0.001** |
| | Asymptote coefficient | 0.035 | -0.062 | 0.136 | 1000 | 0.506 |
| Bottom asymptote | Intercept | -0.005 | -0.081 | 0.065 | 1553 | 0.88 |

| | | | | | | |
|---|---|---|---|---|---|---|
| | Species mean body mass | -0.003 | -0.057 | 0.008 | 1000 | 0.668 |
| | Individual body mass | -0.004 | -0.268 | 0.017 | 1000 | 0.71 |
| | $Inflection_{dilution}$ | $-9.6 \times 10^{-4}$ | -0.008 | 0.008 | 1700 | 0.982 |
| | **$Inflection_{antibacterial\ capacity}$** | **-2.87** | **-4.16** | **-1.51** | **1000** | **<0.001** |
| | **Slope** | **$1.2 \times 10^{-4}$** | **$2.66 \times 10^{-5}$** | **$2.31 \times 10^{-4}$** | **1000** | **0.026** |
| | **Top asymptote** | **2.46** | **1.41** | **3.58** | **1000** | **<0.001** |
| | Asymptote coefficient | $9.7 \times 10^{-6}$ | $-1.23 \times 10^{-4}$ | $1.56 \times 10^{-4}$ | 1000 | 0.87 |
| $Inflection_{antibacterial\ capacity}$ | Intercept | -3.54 | -0.083 | 0.008 | 1000 | 0.156 |
| | Species mean body mass | $-8.7 \times 10^{-5}$ | -0.008 | 0.007 | 887.6 | 0.992 |
| | Individual body mass | -0.004 | -0.016 | 0.008 | 1136.3 | 0.544 |
| | **$Inflection_{dilution}$** | **1.17E** | **0.007** | **0.016** | **864.4** | **<0.001** |
| | **Slope** | **$3.5 \times 10^{-4}$** | **$3.0 \times 10^{-4}$** | **$3.9 \times 10^{-4}$** | **1000** | **<0.001** |
| | **Bottom asymptote** | **-8.34** | **-1.22** | **-0.045** | **1000** | **<0.001** |
| | **Top asymptote** | **7.95** | **7.81** | **8.12** | **1000** | **<0.001** |
| | **Asymptote coefficient** | **$-2.27 \times 10^{-4}$** | **$-3.0 \times 10^{-4}$** | **$-1.5 \times 10^{-4}$** | **1000** | **<0.001** |

[a] individual deviation from the species mean

**Table S8.** Posterior means, 95% credible interval (CI), effective sample, and P-values for fixed effect from phylogenetic univariate mixed models. Each model included a single parameter from the antibacterial capacity curve against *M. luteus*. $\text{Inflection}_{\text{antibacterial capacity}}$, top asymptote, and bottom asymptote were +1, $log_{10}$-transformed and body mass and slope were $log_{10}$-transformed prior to analyses

| Response Variable | Fixed effect | Posterior mean | 95% CI Lower | 95% CI Upper | Effective sample | pMCMC |
|---|---|---|---|---|---|---|
| $\text{Inflection}_{\text{dilution}}$ | **Intercept** | **-3.23** | **-4.68** | **-1.82** | **1000** | **<0.001** |
| | Species mean body mass | -0.011 | -0.270 | 0.262 | 1000 | 0.9 |
| | Individual body mass[a] | -0.081 | -0.902 | 0.660 | 1000 | 0.858 |
| | $\text{Inflection}_{\text{antibacterial capacity}}$ | 1.15 | -2.65 | 5.392 | 1000 | 0.572 |
| | Bottom asymptote | 0.504 | -0.337 | 1.39 | 900.3 | 0.272 |
| | **Top asymptote** | **5.63** | **2.65** | **8.83** | **1000** | **<0.001** |
| | Slope | 0.001 | -0.006 | 0.007 | 1000 | 0.878 |
| | Asymptote coefficient | -0.001 | -0.006 | 0.003 | 1000 | 0.526 |
| Top asymptote | **Intercept** | **0.070** | **-0.001** | **0.133** | **1000** | **0.038** |
| | Species mean body mass | -0.003 | -0.015 | 0.009 | 1000 | 0.618 |
| | Individual body mass | 0.002 | -0.015 | 0.022 | 1000 | 0.768 |
| | $\textbf{Inflection}_{\textbf{dilution}}$ | **0.004** | **0.002** | **0.006** | **839.2** | **<0.001** |
| | $\textbf{Inflection}_{\textbf{antibacterial capacity}}$ | **1.18** | **1.15** | **1.22** | **896** | **<0.001** |
| | **Bottom asymptote** | **-0.024** | **-0.051** | **-0.002** | **1000** | **0.05** |
| | **Slope** | **-0.001** | **-0.001** | **-0.001** | **1246** | **<0.001** |
| | **Asymptote coefficient** | $\mathbf{2.8 \times 10^{-4}}$ | $\mathbf{1.7 \times 10^{-4}}$ | $\mathbf{3.9 \times 10^{-4}}$ | **1000** | **<0.001** |
| Slope | **Intercept** | **38.5** | **17.2** | **56.1** | **1000** | **<0.001** |
| | Species mean body mass | -4.30 | -8.339 | -0.301 | 1000 | 0.052 |
| | Individual body mass | 0.959 | -8.869 | 11.429 | 1256 | 0.822 |
| | $\text{Inflection}_{\text{dilution}}$ | 0.054 | -1.159 | 1.026 | 1000 | 0.91 |
| | $\textbf{Inflection}_{\textbf{antibacterial capacity}}$ | **220.7** | **173.8** | **266.7** | **746.1** | **<0.001** |
| | Bottom asymptote | -1.55 | -13.3 | 9.58 | 909.3 | 0.78 |
| | **Top asymptote** | **-178.0** | **-217.7** | **-140.9** | **753.9** | **<0.001** |
| | Asymptote coefficient | -0.007 | -0.070 | 0.050 | 927.3 | 0.802 |

| | | | | | | |
|---|---|---|---|---|---|---|
| Bottom asymptote | Intercept | 0.144 | -0.029 | 0.325 | 1000 | 0.12 |
| | Species mean body mass | -0.029 | -0.062 | 0.008 | 1000 | 0.112 |
| | Individual body mass | -0.043 | -0.109 | 0.027 | 1130.3 | 0.228 |
| | $Inflection_{dilution}$ | 0.004 | -0.004 | 0.011 | 1000 | 0.314 |
| | **$Inflection_{antibacterial\ capacity}$** | **0.398** | **0.076** | **0.797** | **1000** | **0.042** |
| | Slope | $-7.4\times 10^{-5}$ | $-6.6 \times 10^{-4}$ | $4.7 \times 10^{-4}$ | 907.4 | 0.826 |
| | **Top asymptote** | **-0.320** | **-0.605** | **-0.036** | **1000** | **0.034** |
| | **Asymptote coefficient** | **$5.2 \times 10^{-4}$** | **$1.1 \times 10^{-4}$** | **$9.0 \times 10^{-4}$** | **916.4** | **0.008** |
| $Inflection_{antibacterial\ capacity}$ | Intercept | -0.022 | -0.081 | 0.036 | 1000 | 0.478 |
| | Species mean body mass | 0.002 | -0.007 | 0.013 | 1000 | 0.692 |
| | Individual body mass | $4.5 \times 10^{-4}$ | -0.016 | 0.016 | 1000 | 0.928 |
| | $Inflection_{dilution}$ | 0.001 | -0.001 | 0.002 | 1000 | 0.542 |
| | **Slope** | $5.4 \times 10^{-4}$ | $4.2 \times 10^{-4}$ | $6.6 \times 10^{-4}$ | **1000** | **<0.001** |
| | **Bottom asymptote** | **0.020** | **0.003** | **0.039** | **1000** | **0.03** |
| | **Top asymptote** | **0.759** | **0.737** | **0.779** | **1127** | **<0.001** |
| | **Asymptote coefficient** | $-2.5 \times 10^{-4}$ | $-3.5 \times 10^{-4}$ | $-1.6 \times 10^{-4}$ | **1000** | **<0.001** |

**Movie S1.**

Mammals show great diversity in antibacterial dilution curves against *Escherichia coli* (n=111 species). Despite this variation, large mammals have higher antibacterial capacity than small mammals.
